# Metabolic network divergence: polyamine and ethylene dynamics in *Arabidopsis thaliana* and *Solanum lycopersicum*

**DOI:** 10.1101/2025.01.24.634693

**Authors:** Kateřina Cermanová, Petra Bublavá, Mersa Darbandsari, Martin Fellner, Ondřej Novák, Michal Karady

## Abstract

Polyamines are ubiquitously present in all living organisms. In plants, together with phytohormone ethylene, their metabolism plays a crucial role in plant stress and ontogenesis. We have evaluated differences in the responses of model plants *Arabidopsis thaliana* and *Solanum lycopersicum* to abiotic stresses and metabolic modulators based on key metabolite levels. Previous approaches have often focused separately on either polyamines, amino acids, or ethylene precursors. As these pathways are directly interconnected, their simultaneous evaluation can significantly impact our understanding of their core mechanisms.

We have therefore developed a novel and validated liquid chromatography – tandem mass spectrometry based method, enabling quantification of fourteen compounds: the amino acids L-arginine, L-citrulline, L-ornithine; biogenic amines *N*^α^-acetyl-L-ornithine and agmatine; the polyamines putrescine, spermidine, spermine, thermospermine, *N*-acetylputrescine, cadaverine, homospermidine; together with methionine and 1-aminocyclopropane-1-carboxylic acid, serving as key non-volatile precursors of ethylene.

Our analysis revealed distinct metabolic responses between *Arabidopsis* and tomato, highlighted by species-specific differences in polyamine metabolism and ethylene precursors dynamics. Drought and salinity stresses triggered fundamentally different metabolic adjustments, with drought consistently inducing higher metabolite levels and spermine showing stress-specific responses. Metabolic inhibitor treatments with aminoguanidine and L-norvaline revealed further divergencies, mainly demonstrated as significant variations in ethylene precursor levels. For all approaches, *Arabidopsis* displayed more pronounced metabolic fluctuations compared to tomato. These results provide direct insights into contrasting metabolic plasticity and the interconnected roles of polyamines, amino acids, and ethylene precursors in plant responses and adaptations.

## 1. Introduction

Plant adaptation to environmental stresses involves complex regulatory networks and metabolic adjustments, with polyamines and ethylene playing crucial roles as key signaling molecules (Alcázar et al., 2020; Chen et al., 2021). Polyamines (PA) comprise of many different small polycationic molecules ubiquitously present in all living organisms, with putrescine (Put), spermine (Spd), spermidine (Spm) and thermospermine (Tspm) considered to be the major PAs in plant lineages (Blázquez, 2024).

The primary PA biosynthetic route involves amino acids L-arginine (Arg), L-ornithine (Orn) and L-citrulline (Cit). It begins with Arg, which can be converted to Orn via arginase or to biogenic amine agmatine (Agm) via arginine decarboxylase enzyme (ADC; Lou et al., 2020). L-Citrulline, an intermediate in the arginine biosynthetic pathway, serves as an important metabolic connector in these processes (Winter et al., 2015). The regulation of these amino acid intermediates, particularly the arginine-ornithine-citrulline interconversion, plays a crucial role in modulating polyamine levels during stress responses (Majumdar et al., 2016a). L-Ornithine can be directly decarboxylated to Put by ornithine decarboxylase (ODC), while Agm is converted to *N*-carbamoylputrescine and then to putrescine (Blázquez, 2024), through activity of agmatine iminohydrolase (AIH) and *N*-carbamoylputrescine amidohydrolase (NLP1, nitrilase-like protein 1). Spd, Spm and Tspm are then consequently synthesized from Put by addition of an aminopropyl group from decarboxylated *S*-adenosylmethionine (dcSAM). Further modification of these PAs by acetylation adds another layer of PA homeostasis complexity, affecting their transport, degradation and biological functions (Lou et al., 2020). The presence or absence of ODC in different plant species presents an interesting model for studying polyamine biosynthesis regulation. While most plants possess both ADC and ODC pathways, *Arabidopsis thaliana* lacks a functional ODC gene, relying solely on the ADC pathway for putrescine synthesis (Hanfrey et al., 2001). In contrast, tomato (*Solanum lycopersicum*) maintains both pathways (Kwak & Lee, 2001), potentially offering greater metabolic flexibility in polyamine biosynthesis and associated stress responses.

Structurally related to the canonical polyamines, cadaverine (Cad) and homospermidine (Hspd) represent important variants in plant polyamine metabolism. Cadaverine, derived from lysine decarboxylation, shares structural similarities with putrescine and can partially fulfill similar cellular functions (Jancewicz et al., 2016). Homospermidine, synthesized by homospermidine synthase using Put as a substrate, represents an interesting branching point in polyamine metabolism with distinct physiological roles (Niemüller et al., 2012).

The molecular crosstalk between polyamines and ethylene biosynthesis represents a critical regulatory node in plant responses. Abiotic stresses, particularly drought and salinity, significantly impact polyamine metabolism (Blázquez, 2024) and ethylene biosynthesis (Achard et al., 2006). Both pathways share *S*-adenosyl-L-methionine (SAM) as a common precursor, where SAM, synthesized from amino acid L-methionine (Met), can either be directed toward polyamine biosynthesis via *S*-adenosylmethionine decarboxylase (SAMDC) or toward ethylene production through 1-aminocyclopropane-1-carboxylic acid (ACC) synthase (Pattyn et al., 2021). This metabolic competition for SAM creates an intricate regulatory network where increased flux through one pathway can affect the other, creating a competitive or cooperative network (Apelbaum et al., 1981; Bürstenbinder et al., 2010; Harpaz-Saad et al., 2012; Van de Poel et al., 2013; Xuan et al., 2024).

Most of small active compounds in plants tend to have diverse effects at the same stage of development, or multiple compounds might influence a single process at once. Their activity is determined by various factors, with concentration being particularly crucial. Understanding their precise quantities is essential for revealing their mechanisms and impacts on plant life. Their levels can vary widely depending on the plant tissue type, often being present in very low amounts. Thus, an effective measurement method must account for these varying concentrations and address the complexities of plant tissue (Petřík et al., 2024). Previous analytical approaches have often focused separately on either polyamines, amino acids, or ethylene precursors, limiting our understanding of their dynamic interrelationships.

Here, we present a novel and validated liquid chromatography – tandem mass spectrometry (LC-MS/MS) based method enabling simultaneous quantification of fourteen metabolically related compounds: the amino acids L-arginine, L-citrulline, L-ornithine, biogenic amines *N*^α^-acetyl-L-ornithine (AcOrn) and agmatine; polyamines putrescine, spermidine, spermine, thermospermine, *N*-acetylputrescine (AcPut), cadaverine, and homospermidine; and the Yang cycle members L-methionine and ACC (1-aminocyclopropane-1-carboxylic acid). We explore their relationship *in vitro* in model plants *A. thaliana* and *Solanum lycopersicum* (tomato) under various abiotic stresses (salinity and drought) and by employing PA metabolism modulators aminoguanidine (AG) and L-norvaline (Nor). This comprehensive analytical approach allows for precise tracking of metabolic flux through both polyamine biosynthetic pathways and ethylene precursor formation, providing insight into the dynamic relationship between these crucial signaling molecules. Such simultaneous quantification is particularly valuable for understanding metabolic adjustments during stress responses, where rapid changes in multiple pathway intermediates can significantly impact plant adaptation mechanisms.

## 2. Materials and methods

### 2.1 Plant material and growth conditions

All plant experiments were conducted using *Arabidopsis thaliana* (L.) ecotype Columbia-0 (Col-0) and *Solanum lycopersicum* ‘Ailsa Craig’ *in vitro* grown seedlings. *Arabidopsis* seeds were first surface sterilised using 70% ethanol supplemented with 0.1% TWEEN-20 for 5 min and then sown on half-strength solid Murashige and Skoog medium (MS; 2.2 g L^-1^ MS media, 7 g L^-1^ agar, pH 5.7) supplemented with sucrose (10 g L^-1^) in square Petri dishes. Seeds were then stratified for two days in 4 °C and placed vertically in a growth chamber (Microclima 1000E, Snijders Scientific) under long-day conditions (16 h light/8 h dark, 23°C/18°C) with a light intensity of 110 μmol photons m^−2^ s^−1^.

Tomato seeds were soaked in 3% sodium hypochlorite solution for 20 min, followed by washing with sterilized distilled water and then sown on described MS medium. To induce germination, the Petri dishes were placed in growth chamber in the dark. After three days, germinated seeds were chosen and transferred to new plates and further used in experiments.

### 2.2 Inhibitor and stress treatments

For stress experiments, 10 days after germination (DAG) old *Arabidopsis* and 4 DAG old tomato seedlings were carefully transferred to new plates for 48 h stress treatment. For salt stress treatment, sodium chloride (NaCl) was added to MS medium prior to autoclaving at three different concentrations: 10 mmol L^-1^ for low, 150 mmol L^-1^ for moderate and 300 mmol L^-1^ for high stress severity. Drought stress was induced using a modified protocol described by Verslues et al. (2006). Two half strength MS media were prepared (2.2 g L^-1^ MS media, 1.2 g L^-1^ MES, pH 5.7), one supplemented with sucrose (10 g L^-1^) and agar (15 g L^-1^) prior to autoclaving, the other with added polyethylene glycol (PEG) after sterilization by autoclaving. For low, moderate and high stress severity, the water potential of approximately -0.5 MPa (250 g L^-1^ PEG), -0.7 MPa (400 g L^-1^ PEG) and -1.2 MPa (550 g L^-1^ PEG) were chosen, respectively. To ensure the final water potential of the plates for stress treatment, the PEG overlay solution was poured onto solidified MS medium in plates in a 2:3 volume ratio (agar:PEG overlay) and equilibrated in sterile conditions overnight. The excess aqueous solution in plates was then poured out just before the seedlings transfer the following day. For control conditions, seedlings were transferred to MS medium without additional sodium chloride and no changes to water potential.

To further study the homeostasis of polyamines, two different inhibitors of the polyamine biosynthesis pathway were used: L-norvaline and aminoguanidine. 5 DAG old *Arabidopsis* and 4 DAG old tomato seedlings were transferred to new plates containing individual inhibitors at multiple concentrations (0.1 mmol L^-1^, 1 mmol L^-1^ and 10 mmol L^-1^) and harvested at two different timepoints – 48 h and 96 h after transfer.

The seedlings were removed from Petri dishes, placed on black linen cloth and photographed (tomato) or scanned still in Petri dishes before and after each individual treatment using EPSON Perfection V550 photo scanner (*Arabidopsis*). All plant samples were collected as whole seedlings, snap frozen in liquid nitrogen and stored in -80 °C until further use.

### 2.3 Metabolite extraction and LC-MS/MS instrumentation

Plant material stored at -80°C was used to prepare technical replicates of approximately 10 mg by pooling and homogenization of all whole seedlings collected from one treatment. Extraction was carried out by adding 500 μL of cold 10% acetonitrile (ACN) with 0.5% formic acid (FA; v/v) along with a mixture of isotopically labelled internal standards (IS, 30 pmol per sample) and three ceria-stabilized zirconium oxide 2 mm beads to the plant material. Samples were further homogenized on bead mill (Retsch GmbH, Haan, Germany) in pre-cooled clamps for 5 min at 27 Hz, twice, followed by centrifugation at 30 000 g for 10 min at 4 °C. Next, 250 μL of supernatant was transferred to a glass insert in a chromatographic vial and evaporated to dryness on rotary vacuum evaporator. Before LC-MS/MS analysis, analytes were derivatized using the AccQ-Tag Ultra Derivatization Kit from Waters, with solutions prepared according to the manufacturer’s instructions. 14 μL of borate buffer and 4 μL of derivatization reagent solution was added to evaporated sample, left on lab bench for 10 minutes and followed by adding 52 μL of 0.1% FA (v/v).

Quantification of polyamines by liquid chromatography – tandem mass spectrometry was performed on Agilent 6495B Triple-Quad MS coupled to 1260 Infinity II LC system (Agilent Technologies, Inc., Santa Clara, CA, USA). For method development, chromatographic column Kinetex Polar (150 x 2.1 mm, 2.6 µm, Phenomenex) was initially tested and Luna Omega PS C18 100Å (150 x 2.1 mm, 1.6μm, Phenomenex) preheated to 53 °C was finally used for all measurements. The total runtime of one analysis is 25 min. As mobile phases, 0.1% FA in water (A) and 0.1% FA in ACN (B) (both v/v) at flow rate of 0.3 ml/min were used in the following non-linear gradient conditions: 0 min – 99% A, 18 min – 83% A, 19 min – 4% A, 19.4 min – 4% A, 20.5 min – 99% A, 25.0 min – 99% A. The resulting LC-MS/MS chromatogram of examined polyamines and related compounds with their respective retention times is shown in Fig. 1.

**Fig. 1.**
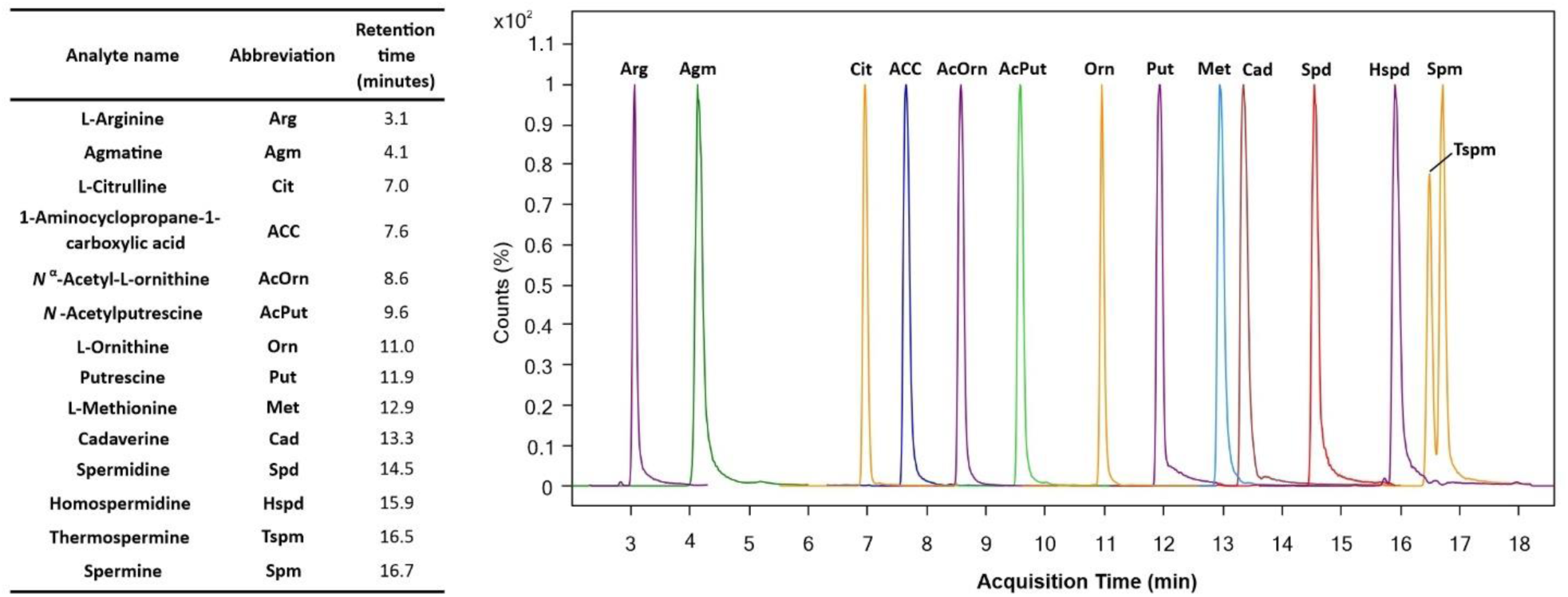
Optimized LC-MS/MS chromatogram of examined polyamines and related compounds. Separation of analytes was achieved using a Luna Omega PS C18 100Å (150 x 2.1 mm, 1.6μm) chromatographic column with 0.1% formic acid in water and with 0.1% formic acid in acetonitrile as mobile phase A and B, respectively. The composition of gradient elution was 0 min – 99% A, 18 min – 83% A, 19 min – 4% A, 19.4 min – 4% A, 20.5 min – 99% A, 25.0 min – 99% A. Measurements were performed on Agilent 6495B Triple-Quad LC/MS coupled to 1260 Infinity II LC system.

The MS source parameters for these analyses were: gas temp 160 °C, gas flow 14 L/min, sheat gas temp 390 °C and shear gas flow 12 L/min. The “Dynamic multiple reaction monitoring (dMRM)” mode and electrospray ionization (ESI) in positive mode were employed with the capillary voltage set to 2800 V and nozzle voltage at constant 0 V.

Quantification of analytes in samples was done using the standard isotope dilution method (Rittenberg and Foster, 1940) with calibration curves prepared using the chemical standards of analytes and the used IS (Supplementary Table S1).

### 2.4 Method validation

Results for method validation are shown in Supplementary Table S2. Our method validation involved evaluating method accuracy and precision, process efficiency, matrix effect, and method recovery, following the guidelines of Matuszewski et al. (2000), using the postextraction addition technique. Sets of 4 replicates of 10 DAG old *Arabidopsis* seedlings of approximately 10 mg were prepared to validate the method. Sample sets were divided into two groups, spiked before extraction and spiked before evaporation, and otherwise processed as described in 2.3. Metabolite extraction and LC-MS/MS instrumentation. The first set of samples was spiked with a known amount of IS before extraction and no authentic standards, other sets were spiked with the same known amount of IS and with non-labelled standards. The spiked content reflects the upper linear range of each analyte (Supplementary Table S1). Due to the broad differences in the abundance levels of the analysed compounds in plants, for validation purposes we decided to include also spiking concentrations reflecting endogenous levels detected in first set of samples (Supplementary Table S2).

Similarly, corresponding sample sets were prepared by spiking samples after extraction and before evaporation with the same amounts of IS and non-labelled standards to calculate method recovery by dividing the response of non-labelled standards from a sample spiked before extraction by the mean response of the corresponding samples spiked before evaporation. The value for each analyte is expressed in percentage.

To determine method accuracy, the endogenous concentration of polyamines from the first non-spiked set of samples was subtracted from the calculated concentrations of spiked sets and compared to the theoretical amount of added non-labelled standards. Results are expressed as %bias.

Process efficiency and matrix effect were expressed as the ratio of the mean of peak areas of analytes spiked before extraction and before evaporation, respectively, and the mean of peak area of the corresponding standard. Finally, the average %RSD values of quantified analytes calculated for all four replicates of each sample set represent the method precision.

We proceeded to evaluate the stability of compounds in stock solutions and plant material, stored in autosampler at 6 °C. Stock solutions were represented by low or high levels of analyte standards with a constant amount of IS mixed together and measured on three consecutive days. The intervals between injections on each day were approximately 24 h. The ratio of analyte and IS peak area was calculated for all compounds and all three timepoints. The resulting values presented as %RSD along with the concentrations of non-labelled standards are displayed as autosampler stability in Supplementary Table S3.

Plant samples stability was evaluated as interday precision, using four replicates of plant samples containing IS in the same intervals and conditions as described above for autosampler stability. Similarly, the resulting values as %RSD are summarized in Supplementary Table S3.

### 2.5 Data processing and statistical analysis

The LC-MS/MS data were processed and quantified using MassHunter Workstation Software ver. B.09.00 (Agilent Technologies). Statistical significance was evaluated using Welch’s one-way ANOVA (p<0.05; p<0.01 and p<0.001). The length of primary and lateral roots was measured using ImageJ v.1.53g (https://imagej.nih.gov/ij/).

## 3. Results

### 3.1 Method development and validation

Proper choice of analytical technique and its variable components is the most important decision of method development; however, the choice mostly reflects the availability of instrumentation at the lab. LC-MS/MS is the most preferred option in phytohormonal and generally small metabolite analysis (Cao & He, 2023). Due to the polar character of polyamines, amino acids and ACC, thus weak retention on classic LC setups with C18 columns, derivatization is commonly used as an approach for these compounds analysis. The main strength of derivatization lies in in its ability to enhance sensitivity of the method, chemical stability and improvement of chromatographic properties (retention) of the compounds, increased specificity and selectivity etc. (Jain & Verma, 2018). Many different approaches have been used for various biogenic amines and polyamines, employing benzoyl chloride or other derivatization reagent (Magnes et al., 2014; Wong et al., 2016; Ćavar Zeljković et al., 2024). Our choice is the robust and tested AccQ-Tag Ultra Derivatization Kit from Waters. It employs 6-aminoquinolyl-*N*-hydroxysuccinimidyl carbamate based derivatization. This kit is convenient because it comes as a ready-to-mix solution, requires minimal effort, and offers a quick (<10 minutes) and reliable reaction executable on lab bench. It produces stable amino acid, primary and secondary amine derivatives, that are ideal for tandem mass spectrometry analysis (Boughton et al., 2011) and it has been used before for subsequent polyamine and amino acids analysis, eg. in human serum (Gray et al., 2017), plants (Boughton et al., 2011; Gaudin et al., 2014) and for ACC quantification from *Arabidopsis* (Karady et al., 2024).

Plant material extraction, leading to the final sample, is probably the most crucial step in method development, as it offers a very wide variability of options, with probably the only limitations presented by preferably using solutions that are able to quench enzyme activity and also are volatile (to prevent analyte turnover and for final compatibility with MS; Petřík et al., 2024). We opted to try out minimal amount of performed steps, which consist of plant material extraction/homogenization by addition of extraction solvent, internal standards and zirconium beads, followed by centrifugation to remove solid particles and consequent evaporation of the obtained supernatant to dryness. Final steps involve sample derivatization, followed by derivatization reaction termination by abrupt pH change (see chapter 2.3 Metabolite extraction and LC-MS/MS instrumentation). However, as this approach uses a rather mild extraction solution (10% ACN with 0.5% FA; Karady et al., 2024), our method most likely does not differentiate between free, bound or conjugated polyamines, amino acids and ACC. The main advantages of our method, besides its simplicity of extraction, is the usage of internal standards for most of our analyte content estimation, as well as joining the quantification of polyamine, L-methionine and ACC (as direct ethylene precursor) together in one validated method.

The process of selecting the proper chromatographic column relied on its ability to separate spermine and thermospermine. Although the Kinetex Polar column offers very good partitioning of some *cis-* and *trans-* phytohormone isomers (Karady et al., 2024), a good separation was finally achieved with Luna Omega PS C18 column (see chapter 2.3 Metabolite extraction and LC-MS/MS instrumentation). Although the Tspm and Spm peaks did not achieve baseline separation, their validation procedure was unaffected, with both compounds passing. Upon settling on derivatization, extraction method and following the choice of proper chromatographic column, we proceeded with method validation. Method accuracy and precision, process efficiency, matrix effect, method recovery with autosampler stability and interday precision were assessed for all compounds (see chapter 2.4 Method validation). For all examined analytes, values for method accuracy and precision, autosampler stability and interday precision stay on average below 15%. The lowest process efficiency was calculated for Agm (7.6%), the highest for Hspd (169.6%), average matrix effect similarly varied for different analytes (10.6% for ACC, 248.2% for Arg). Finally, average method recovery was estimated to be the lowest for Orn (78.6%) and the highest for Cit (210.7%; Supplementary Table S2).

### 3.2 Metabolite and phenotypical changes under salt and drought stress in *A. thaliana* and tomato

Salt and drought stress experiments were performed by transferring 10 (*Arabidopsis*) or 4 (tomato) DAG old plants for 48 h treatment by 10, 150 and 300 mM NaCl concentration and a changing water potential (-0.5; -0.7 and -1.2 MPa). Morphological changes are shown in Supplementary Fig. S1 for salinity and in Supplementary Fig. S2 for drought stress. For both stresses and both plants, primary root (PR) growth was significantly inhibited only at highest stress severity. Metabolite changes for both stresses and both plants are shown in Fig. 2 and Fig. 3. In *Arabidopsis* salinity stress, ACC content at 150 mM has weak correlation with Met. In tomato ACC has a dramatic increase in the highest salt concentration. This might point to a fact that 300 mM NaCl concentration represents a severe salinity stress, where prolonged exposure does not reflect only adaptation changes, but may result in plasmolysis of root cells and salt shock instead of stress (Shavrukov, 2013). In *Arabidopsis*, polyamines and amino acids generally decrease in comparison to control conditions - at 300 mM, a complete depletion of Cad and most significant changes for almost all of the compounds are observed. In tomato, the levels seem to be more stable, with Spm showing gradual increase until 150 mM and Agm, Orn, AcOrn, AcPut and Spd having minimal changes. A significant decrease of Put occurs at lower concentration in *Arabidopsis*, compared to tomato.

**Fig. 2.**
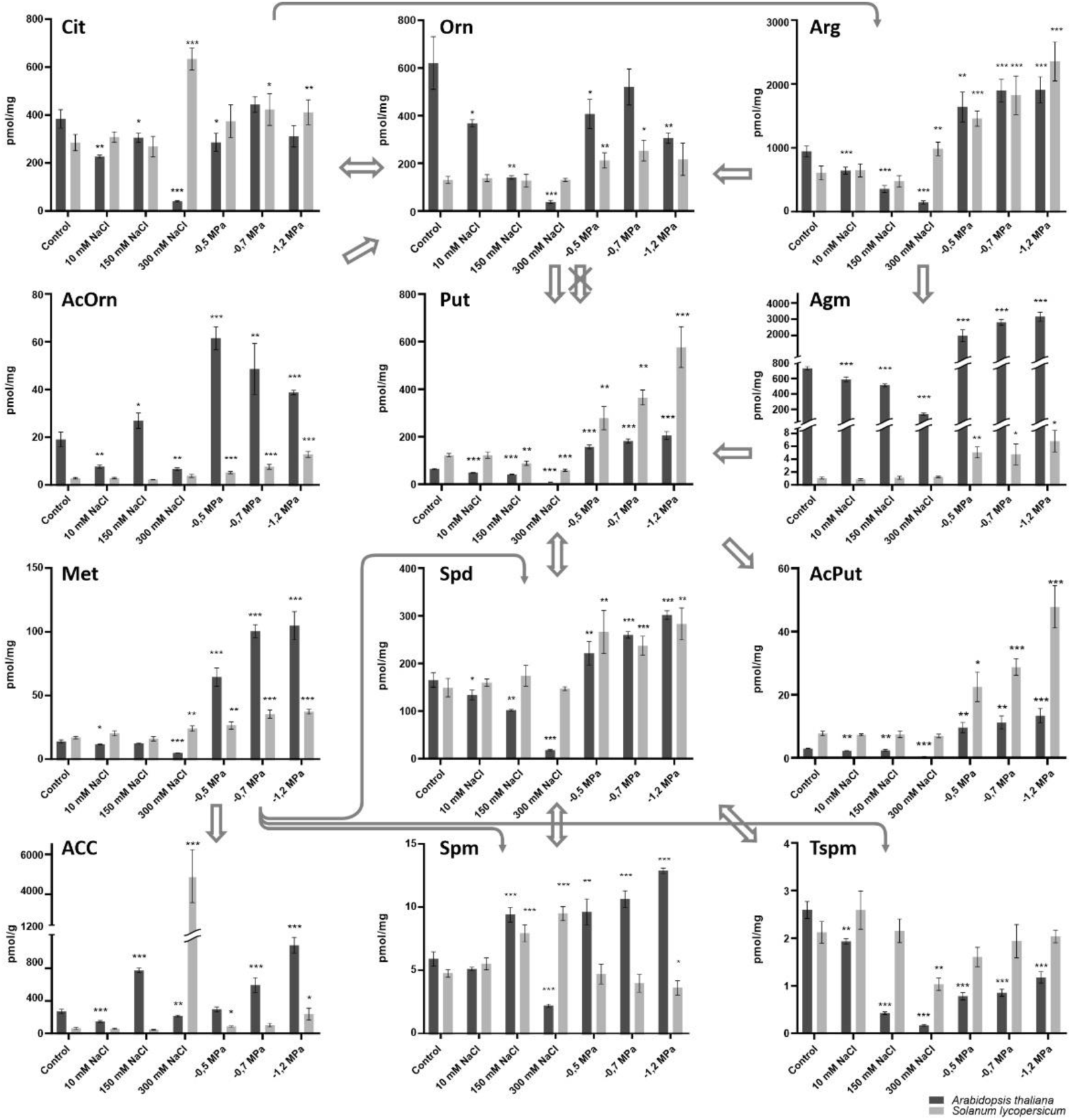
Polyamine and related compounds levels in salt and drought stressed *A*. *thaliana* and *S*. *lycopersicum*. Metabolite profiles of polyamines and related compounds in salt and drought stressed 12 DAG old seedlings of *A*. *thaliana* (dark bars) and 6 DAG old seedlings of *S*. *lycopersicum* (light bars) stressed for 48 h. The major polyamines Put, Spd, Spm and Tspm content is depicted along with related amino acids and biogenic amines (Cit, Arg, Orn, Agm), metabolite forms (AcOrn, AcPut) and Yang cycle metabolites (Met, ACC). Stress treatments were done at three levels each, low (10 mM NaCl, -0.5 MPa), moderate (150 mM NaCl, -0.7 MPa) and severe (300 mM NaCl, -1.2 MPa). Metabolite concentrations calculated from 4-5 technical replicates are expressed in pmol/g FW (ACC) and in pmol/mg FW for all other analytes. Error bars represent SD, the number of asterisks indicates significance level (* = p<0.05; ** = p<0.01; *** = p<0.001) in comparison to control, non-stressed plants, using Welch’s ANOVA Test. The arrows represent metabolic fluxes between the compounds, crossed out arrow indicates absence of the responsible enzyme in *Arabidopsis*, based on Lou et al., 2020. ACC, 1-aminocyclopropane-1-carboxylic acid; AcOrn, *N*^α^-acetyl-L-ornithine; AcPut, *N*-acetylputrescine; Agm, agmatine; Arg, L-arginine; Cit, L-citrulline; Met, methionine; Orn, L-ornithine; Put, putrescine; Spd, spermidine; Spm, spermine; Tspm, thermospermine; DAG, days after germination; FW, fresh weight; SD, standard deviation.

**Fig. 3.**
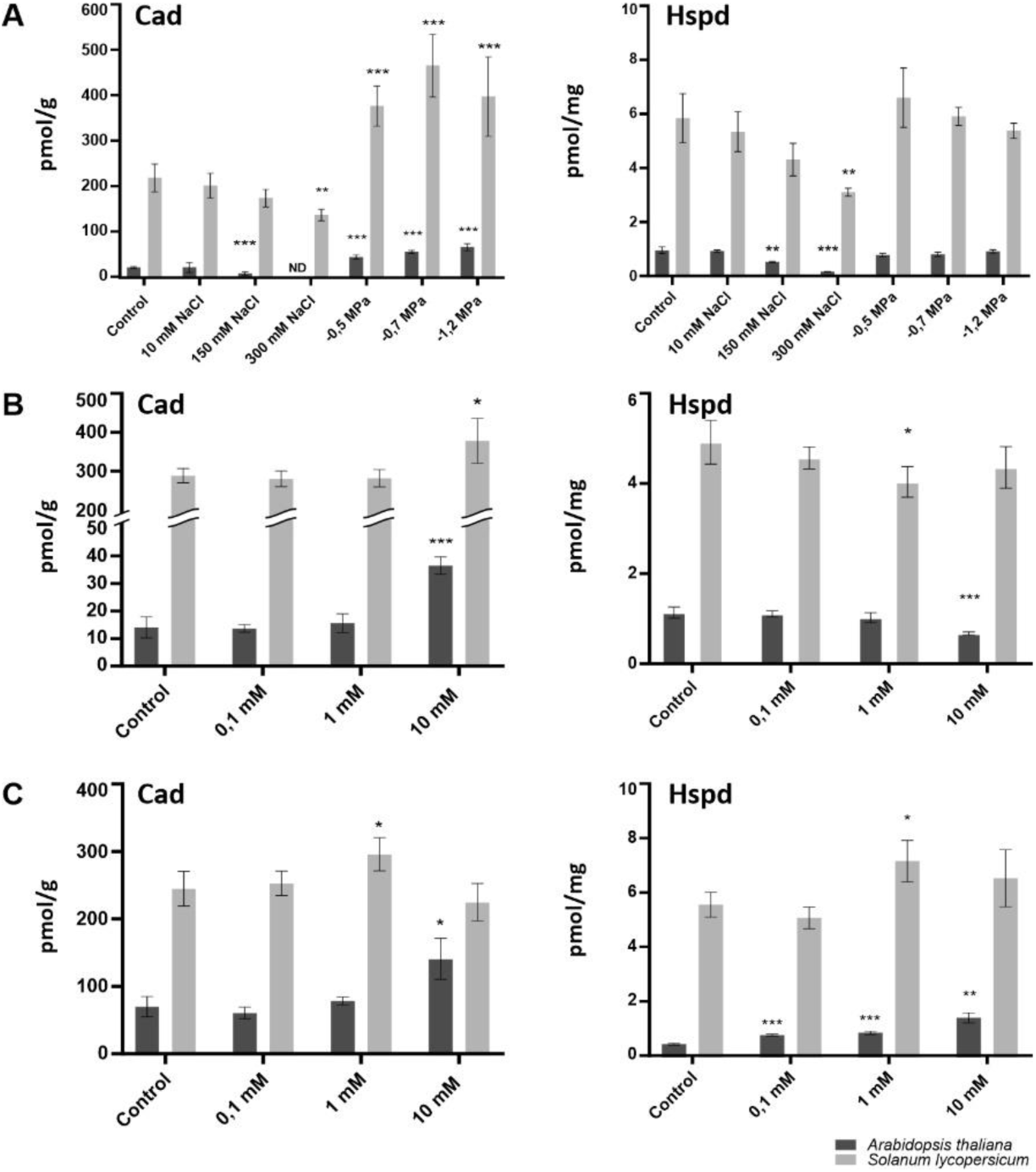
Cadaverine and homospermidine levels in stressed, aminoguanidine and L-norvaline treated *A*. *thaliana* and *S*. *lycopersicum* after 48 h of treatment. (A) Endogenous levels of cadaverine (Cad) and homospermidine (Hspd) in salt and drought stressed 12 DAG old seedlings of *A*. *thaliana* (dark bars) and 6 DAG old seedlings of *S*. *lycopersicum* (light bars). Stress treatments were done for 48 h at three levels each, low (10 mM NaCl, -0.5 MPa), moderate (150 mM NaCl, -0.7 MPa) and severe (300 mM NaCl, -1.2 MPa). (B) Metabolite profiles of Cad and Hspd in 7 DAG old *A*. *thaliana* (dark bars) and 6 DAG old *S*. *lycopersicum* (light bars) seedlings treated with aminoguanidine (AG) for 48 h at three concentration levels, 0.1 mM, 1 mM and 10 mM. (C) Cad and Hspd levels in 7 DAG old *A*. *thaliana* (dark bars) and 6 DAG old *S*. *lycopersicum* (light bars) seedlings treated with L-norvaline (Nor) for 48 h at three concentration levels, 0.1 mM, 1 mM and 10 mM. Metabolite concentrations calculated from 4-5 technical replicates are expressed in pmol/g FW for Cad and in pmol/mg FW for Hspd. Error bars represent SD, the number of asterisks indicates significance level (* = p<0.05; ** = p<0.01; *** = p<0.001) in comparison to control, non-treated plants, using Welch’s ANOVA Test. DAG, days after germination; FW, fresh weight; ND, not detected; SD, standard deviation.

In both species, for drought stress, most of the compounds are gradually elevated in response to increasing stress severity, Cit and Orn display a similar trend. AcOrn and Spm exhibit opposite trajectory (Spm increases in *Arabidopsis*, decreases in tomato), Hspd levels are unaffected, although its precursor, Put, undergoes a significant increase. Spd is also significantly elevated. For Tspm, both plants show initially significantly lower levels than in control. Generally, tomato shows more consistent progressive increases and has better metabolic stability, *Arabidopsis* plants display more significant fold changes. For both species, cadaverine levels are markedly elevated.

### 3.3 Metabolite and phenotypical changes after aminoguanidine treatment in *A. thaliana* and tomato

Aminoguanidine effect on examined compounds pools were performed at three differing concentrations (0.1; 1 and 10 mM), with plants collected after 48 and 96 h of treatment. Results for LC-MS/MS measurements appear in Fig. 4, Fig. 3 (48 h) and Supplementary Fig. S3 and Supplementary Table S4 (96 h). Phenotypical changes are displayed in Fig. 5 (48 h) and Supplementary Fig. S4 (96 h). Phenotypically, AG inhibits PR length and number of lateral roots (LR), with significant effects at its highest, 10 mM concentration. Accordingly, in tomato, changes in PR length and number of LR can be observed only at 10 mM AG, with mild inhibition at 48 h and much more pronounced inhibitory effect at 96 h. In *Arabidopsis*, the number of LR is affected in 1 and 10 mM concentration after 48 h, persisting only for highest concentration after 96 h.

**Fig. 4.**
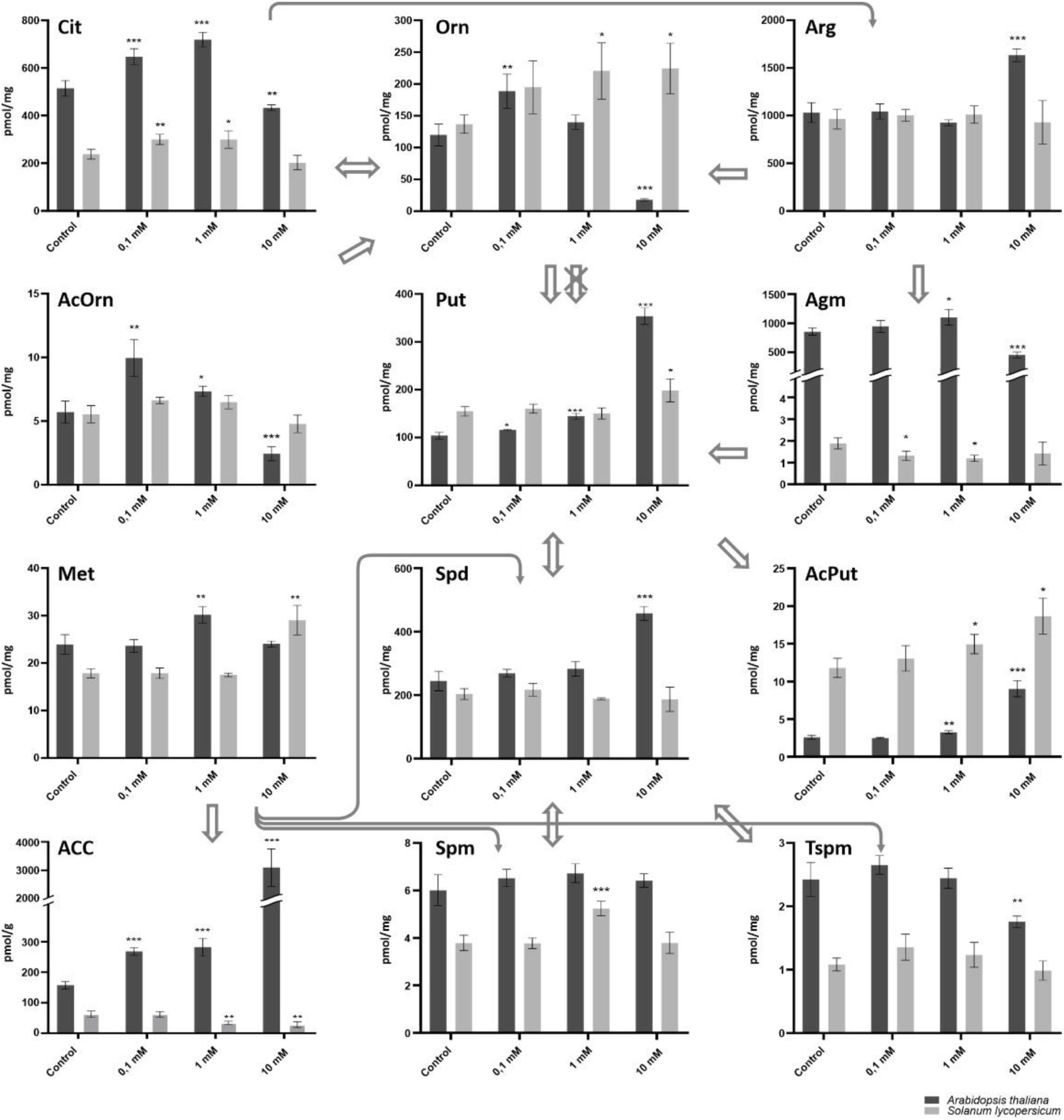
Polyamine and related compounds levels in aminoguanidine treated *A*. *thaliana* and *S*. *lycopersicum* after 48 h of treatment. Metabolite profiles of polyamines and related compounds of 7 DAG old *A*. *thaliana* (dark bars) and 6 DAG old *S*. *lycopersicum* (light bars) seedlings treated with aminoguanidine (AG) for 48 h at three concentration levels, 0.1 mM, 1 mM and 10 mM. The major polyamines Put, Spd, Spm and Tspm content is depicted along with related amino acids and biogenic amines (Cit, Arg, Orn, Agm), metabolite forms (AcOrn, AcPut) and Yang cycle metabolites (Met, ACC). Metabolite concentrations calculated from 4-5 technical replicates are expressed in pmol/g FW (ACC) and in pmol/mg FW for all other analytes. Error bars represent SD, the number of asterisks indicates significance level (* = p<0.05; ** = p<0.01; *** = p<0.001) in comparison to control, non-treated plants, using Welch’s ANOVA Test. The arrows represent metabolic fluxes between the compounds, crossed out arrow indicates absence of the responsible enzyme in *Arabidopsis*, based on Lou et al., 2020. ACC, 1-aminocyclopropane-1-carboxylic acid; AcOrn, *N*^α^-acetyl-L-ornithine; AcPut, *N*-acetylputrescine; Agm, agmatine; Arg, L-arginine; Cit, L-citrulline; Met, methionine; Orn, L-ornithine; Put, putrescine; Spd, spermidine; Spm, spermine; Tspm, thermospermine; DAG, days after germination; FW, fresh weight; SD, standard deviation.

**Fig. 5.**
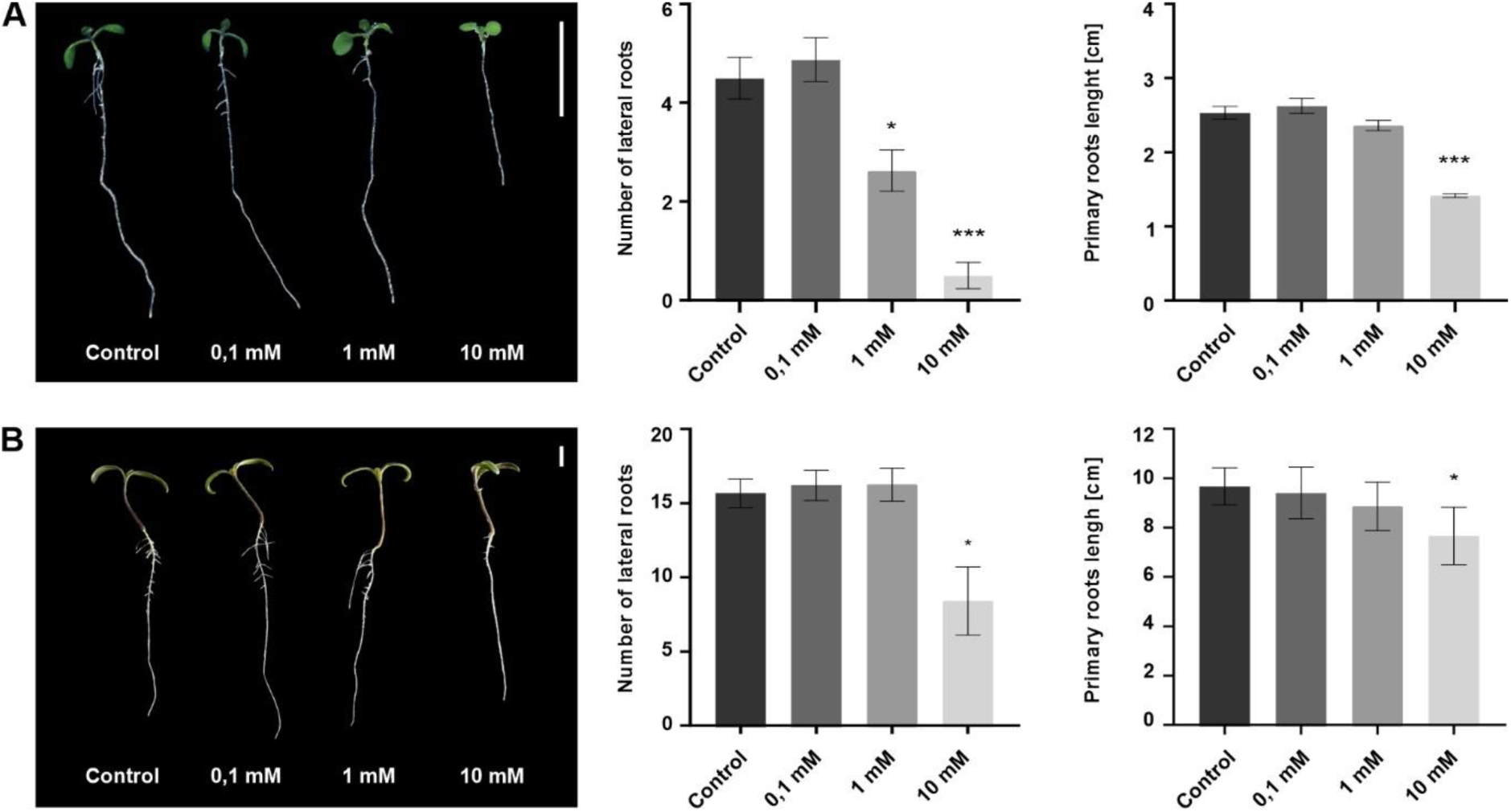
Phenotypes of aminoguanidine treated *A*. *thaliana* and *S*. *lycopersicum* after 48 h of treatment. Phenotype of 7 DAG old *A*. *thaliana* (A) and 6 DAG old *S*. *lycopersicum* (B) seedlings treated with aminoguanidine (AG) for 48 h at three concentration levels, 0.1 mM, 1 mM and 10 mM. Comparison of number of lateral roots of seedlings (central) and the length of their primary roots (right) measured using ImageJ software. Values were calculated from 4-8 seedlings. Error bars represent SE, the number of asterisks indicates significance level (* = p<0.05; ** = p<0.01; *** = p<0.001) in comparison to control, non-treated plants, using Welch’s ANOVA Test. Scale bar = 10 mm; SE, standard error.

For metabolic data, ACC response is significantly opposite in tomato and *Arabidopsis* after 48 h, with content stabilization after 96 h, except for the highest AG concentration, where the opposite trend remains and is not reflected in Met concentrations. In both species, Put (especially at highest AG concentration) is significantly elevated after 96 h with more stable content in tomato, Spm levels remain fixed (with the only exception in 1 mM treatment for tomato after 48 h) and Spd is significantly affected only at highest AG concentration for *Arabidopsis*. Arg and AcOrn content remains stable in tomato after 48 and 96 h. Tspm increases only after 96 h treatment. For *Arabidopsis*, Arg and AcPut follow the same pattern. Tspm levels are influenced only at highest AG concentration for both time points.

### 3.4 Metabolite and phenotypical changes after L-norvaline treatment in *A. thaliana* and tomato

For L-norvaline, identically to AG treatment, two different time-points were used to collect control plants and plants treated with three different Nor concentrations (0.1; 1 and 10 mM). Phenotype differences (Fig.6 for 48 h; Supplementary Fig. S5 for 96 h) display much more pronounced effects for *Arabidopsis*. Both PR length and LR density is robustly negatively affected at all treatments. Tomato phenotype is much more resilient, with inhibitory effects observed only at highest concentrations, and a mild influence on PR length at second highest concentration after 96 h.

**Fig. 6.**
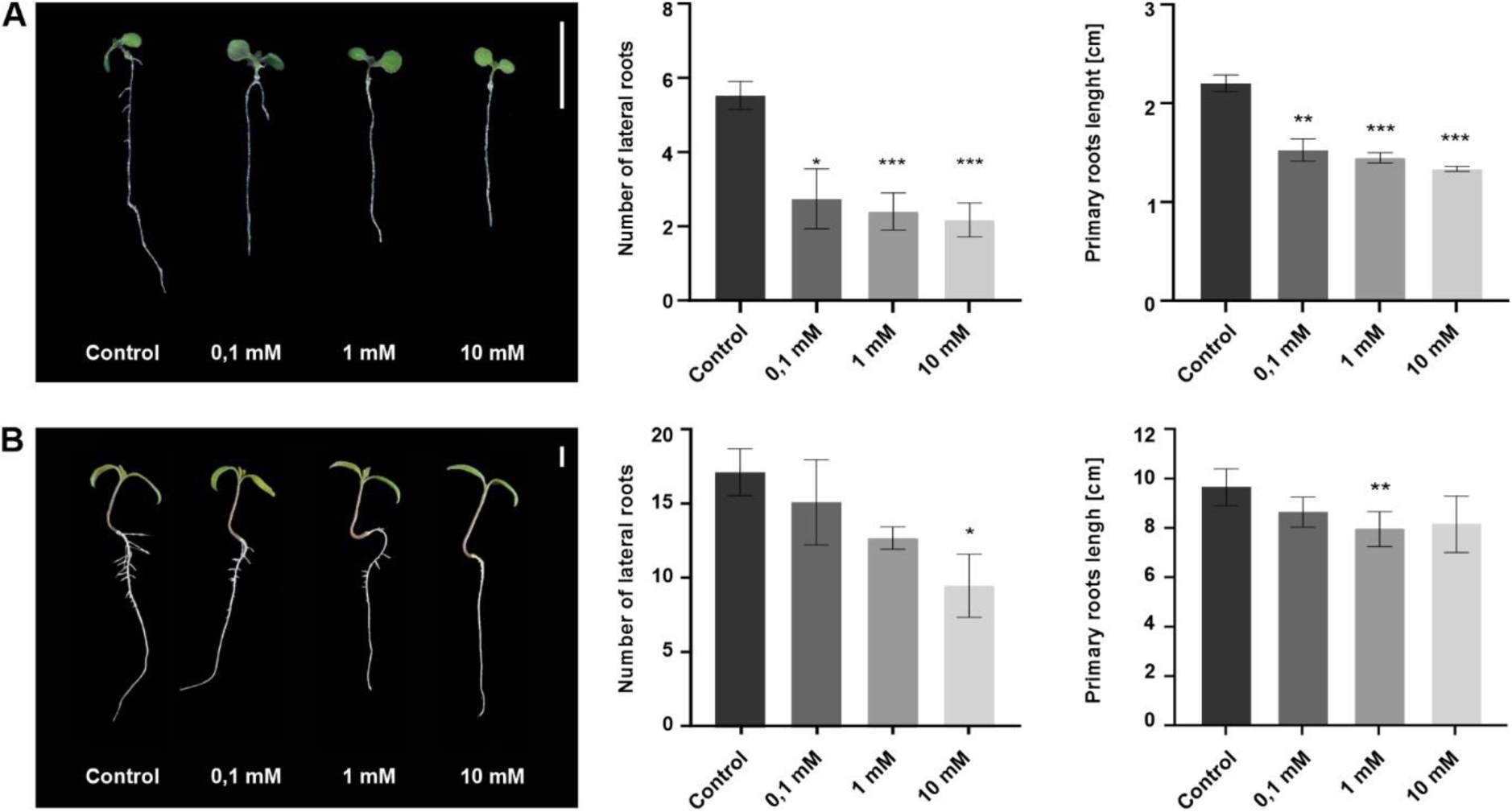
Phenotypes of L-norvaline treated *A*. *thaliana* and *S*. *lycopersicum* after 48 h of treatment. Phenotype of 7 DAG old *A*. *thaliana* (A) and 6 DAG old *S*. *lycopersicum* (B) seedlings treated with L-norvaline (Nor) for 48 h at three concentration levels, 0.1 mM, 1 mM and 10 mM. Comparison of number of lateral roots of seedlings (central) and the length of their primary roots (right) measured using ImageJ software. Values calculated from 4-8 seedlings. Error bars represent SE, the number of asterisks indicates significance level (* = p<0.05; ** = p<0.01; *** = p<0.001) in comparison to control, non-treated plants, using Welch’s ANOVA Test. Scale bar = 10 mm; SE, standard error.

Compounds measurements are displayed in Fig. 7, Fig.3 for 48 h treatment. The results of 96 h exposure can be found in Supplementary Fig. S6 and Supplementary Table S4. ACC shows a steady increase, with massive spike under highest treatment concentration for both species, with the same trend in Orn concentration. Arg level is initially stable in *Arabidopsis*, with significant uprise after 96 h. For tomato, Arg content goes up after 48 h and gets more stable after 96 h – opposite to *Arabidopsis* results. Cit, Met and Put, AcOrn and AcPut levels are upregulated in *A. thaliana* and tomato. Spd, Spm and Tspm levels are very stable after 48 h, with Tspm remaining unchanged after 96 h, whereas Spd and Spm levels show a contrasting pattern, with both compounds levels raising in *Arabidopsis* and getting lower in tomato. Agm is only weakly affected in *Arabidopsis* but generally going up, in opposition with tomato levels.

**Fig. 7.**
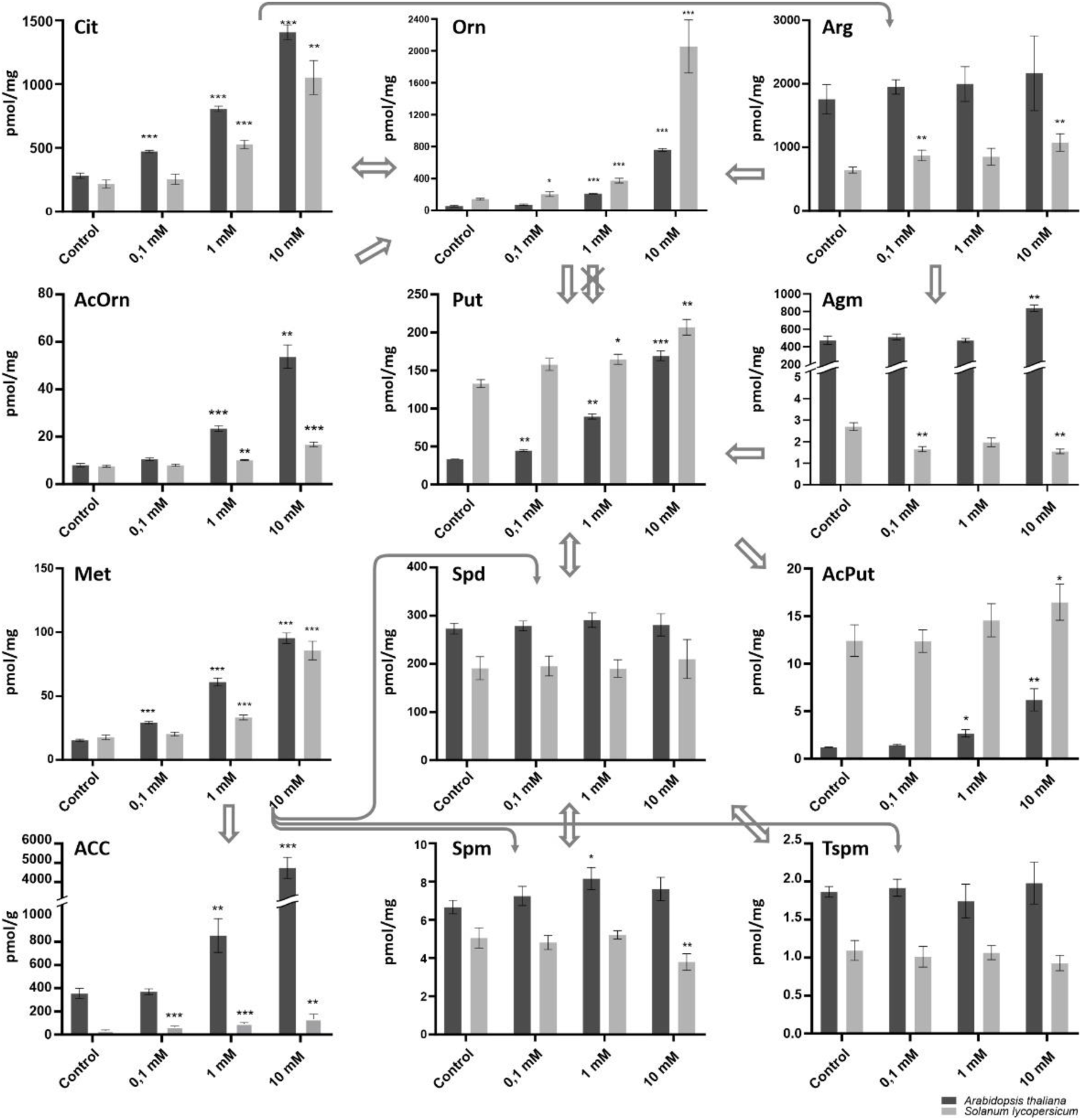
Polyamine and related compounds levels in L-norvaline treated *A*. *thaliana* and *S*. *lycopersicum* after 48 h of treatment. Metabolite profiles of polyamines and related compounds of 7 DAG old *A*. *thaliana* (dark bars) and 6 DAG old *S*. *lycopersicum* (light bars) seedlings treated with L-norvaline (Nor) for 48 h at three concentration levels, 0.1 mM, 1 mM and 10 mM. The major polyamines Put, Spd, Spm and Tspm content is depicted along with related amino acids and biogenic amines (Cit, Arg, Orn, Agm), metabolite forms (AcOrn, AcPut) and Yang cycle metabolites (Met, ACC). Metabolite concentrations calculated from 4-5 technical replicates are expressed in pmol/g FW (ACC) and in pmol/mg FW for all other analytes. Error bars represent SD, the number of asterisks indicates significance level (* = p<0.05; ** = p<0.01; *** = p<0.001) in comparison to control, non-treated plants, using Welch’s ANOVA Test. The arrows represent metabolic fluxes between the compounds, crossed out arrow indicates absence of the responsible enzyme in *Arabidopsis*, based on Lou et al., 2020. ACC, 1-aminocyclopropane-1-carboxylic acid; AcOrn, N^α^-acetyl-L-ornithine; AcPut, N-acetylputrescine; Agm, agmatine; Arg, L-arginine; Cit, L-citrulline; Met, methionine; Orn, L-ornithine; Put, putrescine; Spd, spermidine; Spm, spermine; Tspm, thermospermine; DAG, days after germination; FW, fresh weight; SD, standard deviation.

## 4. Discussion

The transition from method validation to practical application is a critical step, driven by the need to accurately monitor and understand the metabolic profiles of *A. thaliana* and tomato. This progression ensures that our validated method is not only theoretically sound, but also practically effective in real-world scenarios. By proceeding with the measurement in *Arabidopsis* and tomato, we aim to generate comprehensive data that will enhance our understanding of polyamine and related compounds metabolism, their role in plant physiology and relation to ethylene metabolism. It needs to be taken in consideration that our method does not distinguish between bound (insoluble), conjugated (soluble), or free compounds (Chen et al., 2019). As our extraction solution does not match the parameters, such as strong acidity etc., of solutions typically used for release or hydrolysis of conjugated and bound compounds (eg. Roussos & Pontikis, 2007), the results we present are most likely representative of free compounds. A summary of all LC-MS/MS measured contents, for all treatments, in pmol/g or pmol/mg FW, can be found in Supplementary Table S4. Our method encompasses all the analytes present in recently published metabolic pathway of polyamines and related compounds (Lou et al., 2020), with exception of *N*-carbamoylputrescine, *N*^δ^-acetylornithine and glutamate-5-semialdehyde, however with addition of ACC and Met as members of the adjacent Yang cycle. Cit and Orn are presumed to be precursors of Arg as a part of urea cycle (Slocum, 2005). Met precedes the biosynthesis of tri-, tetrapolyamines (Spd, Spm and Tspm) from decarboxylated *S*-adenosylmethionine (Blázquez, 2024); and ACC as a direct non-volatile precursor of the indispensable plant hormone ethylene (Pattyn et al., 2021).

The presented data bring an insight into these compounds average levels in *in vitro* grown tomato and *Arabidopsis* whole seedlings. When comparing data from previous studies with our findings, it is important to note that levels were mostly measured in different tissues, resulting from changing growth conditions, at various stages of plant development and with different methods, extraction solutions, fresh weight versus dry weight etc., which complicates direct comparisons. Our measured content is reported in this paragraph as circa (c.) value, which represents an average of compound levels in control plants from all experiments. Levels of measured ACC (c. 200 pmol/g) for *Arabidopsis* correspond well to previously published results (Cao et al., 2024; Karady et al., 2024; Ziegler et al., 2014), whereas we estimated tomato ACC content to be lower (c. 70 pmol/g) than in *Arabidopsis*. Most ACC levels for tomato are reported from fruit, eg. 50 pmol/g and 2550 pmol/g for unripe and ripe tomato fruit (Bulens et al., 2011), with levels reported for older tomato plants, grown in hydroponic conditions, approximately eight times higher (Zapata et al., 2007). Cadaverine is presumed to be present mostly bound or in its conjugated form (Jancewicz et al., 2016), with reported overall concentrations (5-30 nmol/g FW) being higher than our measurement (c. 33 pmol/g for *A. thaliana* and c. 300 pmol/g for tomato). ACC and Cad were the least abundant compounds in both plants examined by our method. On contrary, Arg showed to be the most abundant analyte in both *A. thaliana* and tomato (c. 1000 and 800 pmol/mg). Previous measurements report Arg to have a bit higher (Frémont et al., 2013) or a bit lower (Modolo et al., 2006) levels in *A. thaliana*, and approximately the same levels in tomato (Kalamaki et al., 2009). Arg takes part in many different and complex processes such as nitrogen flux and storage, however most importantly it is a proteinogenic amino acid (Winter et al., 2015). For L-methionine, our results are c. 17 pmol/mg for *Arabidopsis* with basically the same amount in tomato. Very similar levels have been shown for *Arabidopsis* (Shin et al., 2023) and lower in leaves (Joshi & Jander, 2009). For tomato, similarly to ACC, L-methionine levels are reported mostly for fruit, with levels matching our results (Katz et al., 2006). Met, being also a proteinogenic amino acid as Arg, is known to have lower free levels and lower content in proteome (Hildebrandt et al., 2015). Agmatine amount, in our measurements, shows the biggest differences between *Arabidopsis* and tomato, as we report c. 650 pmol/mg versus a ∼200 fold lower levels of c. 2.7 pmol/mg, respectively. A lower difference (c. 10x) with similar levels in *Arabidopsis* and higher levels for tomato has been reported (Ćavar Zeljković et al., 2024) and slightly lower levels of agmatine have been shown in Jubault et al. (2008) for *Arabidopsis*. In plants, Agm might be prominently present in phenolamide bound forms, eg. p-coumaroylagmatine and other conjugates, having important functions during biotic and abiotic stresses. The substantial difference found in our measurement might be therefore attributed to the varying amounts of p-coumaroylagmatine (or other conjugates), whose contents have been shown to differ significantly among different plant species (Roumani et al., 2021), however the reported amounts for *Arabidopsis* are not that high (Muroi et al., 2009). Another reason for this difference might be the absence of ODC enzyme in *A. thaliana,* as this might lead to a consequence of requiring higher levels of Agm to serve as a biosynthetic pool of Put (and consequently Spd, Spm and Tspm) precursor.

From the examined non-proteinogenic amino acids, L-citrulline and L-ornithine, Cit pools seem to be about two times higher than Orn in both species. Cit levels in our work show very similar levels in *Arabidopsis* and tomato, c. 330 and 250 pmol/mg. Approx. 50 pmol/mg in FW content in *Arabidopsis* has been shown by Planchais et al. (2014). L-Ornithine seems to be an abundant member of polyamine metabolism, with similar amounts in both species (180 and 140 pmol/mg for *A. thaliana* and tomato). Related amounts for *Arabidopsis* (c. 0.4 µmol/g FW) were shown in Majumdar et al. (2016b) and a bit higher amounts (5 µg/100 mg) in Kalamaki et al. (2009). Lower levels for both species were shown in Ćavar Zeljković et al. (2024). *N*^α^-acetyl-L-ornithine data reveals approximately the same concentration in both species (c. 8 pmol/mg for *Arabidopsis* and 6 pmol/mg in tomato), with very similar results obtained for *A. thaliana* in different work (Molesini et al., 2015). Homospermidine is rather rarely measured, we report its levels to be c. 0.8 pmol/mg for *A. thaliana* and higher levels, c. 6 pmol/mg, in tomato. The amount of putrescine in *Arabidopsis* (c. 70 pmol/mg) fits well with other reported content (Majumdar et al., 2017; Molesini et al., 2015). Higher amount was found by us in tomato (c. 150 pmol/mg), comparable with the results of c. 50 nmol/g FW (Jahan et al., 2022) and c. 100-200 nmol/g (D’Incà et al., 2024), but differing with others, reporting lower levels (González-Hernández et al., 2022). *N*-acetylputrescine levels correlate with published literature for both *A. thaliana* and tomato (Ćavar Zeljković et al., 2024) or are a bit lower than previously published in *Arabidopsis* (Lou et al., 2016). Spermidine content is very similar in both tested plants, c. 220 pmol/mg for *Arabidopsis* and c. 205 pmol/mg for tomato. For *Arabidopsis*, comparable numbers were obtained from Li et al. (2023) and Majumdar et al. (2017), while in tomato similar content was reported in stems (D’Incà et al., 2024) and in fruit (Mehta et al., 2002). Spermine levels are the least abundant from the three most studied polyamines (Put, Spd, Spm), with c. 6 and 4.4 pmol/mg for *Arabidopsis* and tomato. Content in tomato fruit is very close to our reported numbers (Mehta et al., 2002) and a bit higher in tomato stems and leaves (D’Incà et al., 2024). In *Arabidopsis*, the reported amount aligns with different published results (Alcázar et al., 2005; Cuevas et al., 2008) whereas higher levels are reported in Majumdar et al. (2017) and Molesini et al. (2015). In this study, thermospermine was found in both species with almost identical levels of c. 2 pmol/mg. Very similar numbers were obtained before for *Arabidopsis* (Naka et al., 2010; Solé-Gil et al., 2019), and for tomato as well (D’Incà et al., 2024).

The content of presented compounds was further examined under various stresses and by treatment with L-Norvaline and aminoguaidine, as enzyme activity modulators. Under salinity stress it needs to be pointed out, that the pronounced effects under the highest concentration (300 mM), where significant swings or reversed trends of some compounds amount are observed, should be considered to be rather a salinity shock than stress (Shavrukov, 2013), as mentioned before, and thus involve multimodal changes that are more severe and separated from homeostasis connected to stress mitigation. We report that polyamines and related amino acids generally decrease after 48 h of salt stress, with the exception of Spm which is elevated in both species, except for 300mM in *Arabidopsis*, where it suddenly decreases, probably as a result of severely depleted Put and Spd (Kasinathan & Wingler, 2004). This corroborates well with Spm taking a special role during abiotic stresses, when compared to other members of polyamine metabolism (Seifi & Shelp, 2019; Yamaguchi et al., 2006). Spd, Agm, AcOrn and AcPut remain stable in tomato. Generally, polyamine supplementation or polyamine biosynthetic genes overexpression alleviates salt stress, with other studies reporting various effects on polyamine levels (Blázquez, 2024). Amino acid levels also decrease in our study for *Arabidopsis,* in contrast with tomato, where the levels remain unaffected for Orn, Agm and, with the exception of the 300 mM treatment, for Arg, Cit and Met. The overall decreasing levels of polyamines are most likely due to salinity-linked activation of PAO (polyamine oxidase), DAO (diamine oxidase) and ADC enzymes (Raziq et al., 2022), probably with combination of phenolamide linked conjugation (Yang et al., 2024). A general decline in *Arabidopsis* PA levels might also be the result of significant Arg reduction, as this simply translates into corresponding Agm, Put and Spd reduction. Setting aside the results of 300 mM treatment, elevated ACC and subsequent ethylene production is a known response to salt stress (Achard et al., 2006), however, we observed stable levels for tomato. Put might also influence ACC levels as it has been shown to elevate ethylene content in salt treated plants (Quinet et al., 2010). In *Arabidopsis*, a decrease and increase was achieved, with the increase at 150 mM fitting well with published data (Achard et al., 2006), however with much less data available for the decrease observed at 10 mM treatment. Tomato ACC and ethylene levels at 100 mM treatment and different conditions, have been shown to increase (Zapata et al., 2007). Hspd follows the pattern of its precursor Put. Cad is stable in tomato and is significantly diminished in *Arabidopsis*.

Drought stress revealed a radically opposite pattern of analyte content compared to salinity stress. Almost all compounds display a significantly upward trend. The employed PEG mediated osmotic stress lacks the ionic component of salinity stress, so the response should be much more water-deficit related, thus simulating drought conditions *in vitro* (Verslues et al., 2006). An upward trend might be simply the result of upward L-arginine trend, inducing increase in polyamine biosynthesis (Hildebrandt, 2018). Most amino acids increase during drought stress, with Arg showing very significant gains (Good & Zaplachinski, 1994). The literature results regarding polyamine content during osmotic stress is rather conflicting. The role of polyamines can vary between different plant species and even among different parts of the same plant, depending on whether they are experiencing osmotic or water stress (Chen et al., 2019). Two compounds displayed opposite trend in *Arabidopsis* vs tomato – AcOrn, Spm and to a degree Orn, pointing to possible activity of ODC or OTC (ornithine transcarbomylase) enzyme under drought stress, especially when compared to salinity stress, where levels of Orn remained unchanged in tomato. Met and ACC are both substantially increased (less in tomato), with Met increase being reflected in Spd and Spm increase in *Arabidopsis* and Spd in tomato. Ethylene biosynthesis is known to be upregulated upon these stress conditions (Khan et al., 2024). Tspm levels stay lower than control in both species and are unaffected in tomato. Cad levels follow the general trend of upregulation in both plants, however, Hspd levels seem to be disconnected from the trends of its precursor Put, by displaying complete stability in both plants.

For both stresses, accumulation of plant stress response osmolytes occurs, with proline being one of the main players (Winter et al., 2015). Feeding plants with Arg or Orn results in elevated proline levels (Liang et al., 2013), therefore this link might also be important.

To further examine the differences between analyte levels in *Arabidopsis* and tomato, we have used two modulators of polyamine and related amino acids metabolism – aminoguanidine and L-norvaline. AG inhibits polyamine (Put, Spm, Spd, Tspm) catabolism and possibly also the nitric oxide synthase, thus affecting urea cycle metabolites/amino acids (Arg, Cit, Orn; Köhler & Szepesi, 2023). Generally, the treatment should lead to elevated Arg, Put and possibly Spd and Spm content. AG presumably has no direct effect on Yang cycle, therefore, we should observe the feedback of modulated polyamine metabolism on ethylene biosynthesis. We report moderate effects of AG treatment. Most profound effect is ACC content upregulation in *Arabidopsis* and downregulation in tomato after 48 h, however, with the effect less present after 96 h, when ACC levels stabilize, but still show a discerning influence at highest treatment concentration. This effect is not reflected in Met levels after 48 h and only marginally after 96 h. ACC content trends in *Arabidopsis* do most closely resemble Put levels, however, influence of Put on ACC levels should be negative, as Put has been shown to inhibit ACC synthase enzyme (Hyodo & Tanaka, 1986). ACC biosynthesis is also modulated by nitric oxide (NO; Jagadis Gupta et al., 2022), whose levels should be affected by AG. With only one exception, levels of Spm are remarkably stable. AG treatment profoundly affected amino acids (Cit, Orn) metabolism in *Arabidopsis*, however, showed limited effect on Arg metabolism in tomato. AG treatment of plants is rare, with other authors obtaining results in rather distinct conditions than ours. For example, Szepesi et al. (2022) report different effect on content of Put, Spd and Spm when under AG, or combined AG and salt treatment in tomato roots. AG has also been reported to stabilize *S*-adenosylmethionine decarboxylase in mammalian cells (Stjernborg & Persson, 1993), which may explain elevated Spd levels, although those may be influenced by elevated Put and Arg. Otherwise, if counting on limited PA catabolism, Put and, to a smaller degree, Spd levels have been affected, with Put being also acetylated to AcPut.

L-norvaline has a very limited number of reports of usage in *A. thaliana* or tomato treatment. In mammalian biology L-norvaline is used mostly to enhance NO production, as, upon treatment, Arg is processed via L-citrulline and nitric oxide route (Ovsepian & O’Leary, 2018). In plant science it is much more often used as an internal standard for other compounds quantification (Urra et al., 2022), as it should not be naturally present in plant tissues. Our stress and AG treatments show that Arg has, naturally, the strongest impact on downstream polyamine levels and enzyme arginase (converting Arg to Orn or Agm to Put) should be considered as the most influential checkpoint in PA metabolism (Köhler & Szepesi, 2023). We therefore decided to, presumably, inhibit Arg conversion into Put and Orn by using L-norvaline as an arginase inhibitor. That way we can examine if PA levels can be directly influenced by other metabolites in presence/absence of ODC. However, in plants, an arginase independent route is present, forming putrescine from Arg through Agm and *N*-carbamoyl-putrescine, which then should reflect the reduction of arginase activity by raising Agm levels. Our results support this conclusion on Agm content only partly, and oppose them for tomato, where Agm levels are reduced (48 h) or intact (96 h). After 48 h, Nor surprisingly raises Put concentration in both plants, and this raise is not reflected in downstream polyamine levels. Tspm levels are completely unaffected in both plants, and after 96 h Spd and Spm display opposite trends in *A. thaliana* (upwards) and tomato (downwards) - despite raised Put levels and almost unchanged AcPut content in tomato. Consequently, Met is significantly upward in both plants, so there should be sufficient amount of SAM for Spd and Spm synthesis. Met content is very well translated into raising content of ACC in *Arabidopsis* and tomato. Orn and Cit are the most affected metabolites in both plants, showing that Nor is most effective in regulating the amino acids of urea cycle. This significant uprise in Orn and Cit is however completely not reflected in PA levels, except Put. In tomato, this points to the activity of ODC enzyme, however in *Arabidopsis*, a different effect, bridging elevated Cit and Orn levels after 48 h into elevated Put, must be active. However, probably even a low, although nonsignificant, elevation in Arg and Agm content in *A. thaliana* might be sufficient to raise Put.

Phenotypes for most of the treatments show reduced PR length and LR number, with more pronounced effect in *Arabidopsis*, in-line with higher tomato resilience observed for compounds profiling. The stunted PR growth and reduced LR could be attributed to the treatments, or either to raised levels of ACC (Lewis et al., 2011; Růžička et al., 2007), with elevated Arg showing similar effect (Funck et al., 2023; Ravelo-Ortega et al., 2021). Polyamines should exert very little effect on PR growth in *A. thaliana* (Tanaka et al., 2019), however, modulation of PA metabolism could lead to changes in hydrogen peroxide levels, which have been shown to alter PR growth (Tsukagoshi et al., 2010).

## 5. Conclusions

Based on our metabolomic analysis of polyamine-related compounds and Yang cycle intermediates in *Arabidopsis* and tomato under salt and drought stress, we can conclude that these stresses trigger fundamentally different metabolic responses. While salt stress generally leads to decreased metabolite levels, drought stress induces an overall increase in both polyamines and Yang cycle members, with generally stronger responses in *Arabidopsis*, suggesting coordinated regulation of these compounds with polyamine metabolism, especially under water-deficit conditions. Certain compounds, such as spermine, show stress-specific responses and species-specific patterns. The presence of ODC enzyme pathway in tomato appears to contribute to better metabolite homeostasis, especially under salt stress, providing additional metabolic flexibility.

The aminoguanidine treatment revealed contrasting responses in polyamine metabolism between *Arabidopsis* and tomato, with *Arabidopsis* showing more severe putrescine accumulation, likely due to its sole reliance on the ADC pathway for its synthesis. The striking ACC accumulation in *Arabidopsis*, coupled with putrescine increase, suggests a previously underappreciated possible connection between polyamine oxidation and ACC metabolism, which surprisingly shows an opposite trend in tomato. Both species demonstrate active Put modification through *N*-acetylation when oxidation is blocked, indicating this as an important alternative metabolic route for polyamine homeostasis. The more moderate response in tomato, including stable spermidine levels and more modest putrescine increases, suggests, that similarly to abiotic stress conditions, availability of ODC provides better homeostatic control of polyamine metabolism.

L-Norvaline treatment significantly impacts polyamine and ethylene precursor metabolism by inhibiting arginase, leading to increased L-arginine, L-citrulline, and L-ornithine levels in both *Arabidopsis* and tomato. This redirection enhances polyamine synthesis, evident from elevated putrescine, spermidine, and spermine levels, with tomato showing higher basal putrescine levels. L-Methionine and ACC also increase, particularly in *Arabidopsis*. The interplay between the Yang cycle and polyamine metabolism suggests that increased L-arginine availability supports both pathways. Tomato exhibits a more balanced response, likely due to dual pathways for polyamine biosynthesis and tighter ethylene regulation.

Overall, these findings reveal the complex and species-specific metabolic responses of *Arabidopsis thaliana* and *Solanum lycopersicum* to abiotic stresses, demonstrating how polyamine and ethylene metabolism is differently modulated under salt and drought conditions, with tomato’s dual biosynthetic pathways providing better metabolic flexibility compared to *Arabidopsis*. The research unveils these metabolic dynamics through a novel LC-MS/MS method that simultaneously quantifies fourteen metabolically interconnected compounds, providing insights into how these plants modulate their metabolism in response to its perturbations.

## Funding

Presented work was supported by The Czech Science Foundation (GAČR) via 20-25948Y junior grant, by Internal Grant Agency of Palacký University (IGA_PrF_2025_019) and by the “Biorefining and circular economy for sustainability” (TN02000044) grant.

## Author contributions

K.C. and M.K. performed and designed the experiments, executed the measurements and wrote the manuscript. P.B. and M.D. carried out sample preparation, collection and statistical analysis. M.F. and O.N. performed research supervision and manuscript review. All authors have revised and agreed to the published version of the manuscript.

## Acknowledgements

The authors thank Hana Svobodová and Veronika Střílková for excellent technical assistance.

## Notes

### Competing Interest Statement

The authors have declared no competing interest.

