## Supplements for "Metabolic network divergence: polyamine and ethylene dynamics in *Arabidopsis thaliana* and *Solanum lycopersicum*"

**Supplementary Material**

List of tables and figures:

**Supplementary Table S1**: Optimized LC-MS/MS parameters for analysed polyamines and related compounds and their corresponding internal standards used for quantification.

**Supplementary Table S2**: Summary of method validation parameters

**Supplementary Table S3**: Autosampler and interday precision of studied polyamines and related compounds, expressed as %RSD.

**Supplementary Table S4**: Examined analyte levels in seedlings of *A*. *thaliana* and *S*. *lycopersicum* under stress conditions and after treatment with aminoguanidine and L-norvaline

**Supplementary Fig. S1**: Phenotypes of *A*. *thaliana* and *S*. *lycopersicum* after 48 h of salinity stress treatment.

**Supplementary Fig. S2**: Phenotypes of *A*. *thaliana* and *S*. *lycopersicum* after 48 h of drought stress treatment.

**Supplementary Fig. S3:** Polyamine and related compounds levels in aminoguanidine treated *A*. *thaliana* and *S*. *lycopersicum* after 96 h of treatment.

**Supplementary Fig. S4:** Phenotypes of aminoguanidine treated *A*. *thaliana* and *S*. *lycopersicum* after 96 h of treatment.

**Supplementary Fig. S5:** Phenotypes of L-norvaline treated *A*. *thaliana* and *S*. *lycopersicum* after 96 h of treatment.

**Supplementary Fig. S6:** Polyamine and related compounds levels in L-norvaline treated *A*. *thaliana* and *S*. *lycopersicum* after 96 h of treatment.

**Supplementary Table S1**: Optimized LC-MS/MS parameters for analysed polyamines and related compounds and their corresponding internal standards used for quantification.

| Analyte | Abbreviation | Internal standard | Precursor ion (*m/z*) | Fragment ion (*m/z*) | Collision energy (eV) | Linear range (pmol) | $R^2$ | Derivatization multiplicity |
|---|---|---|---|---|---|---|---|---|
| **1-Aminocyclopropane-1-carboxylic acid** | **ACC** | **$[^2H_4]$-ACC** | 272.1 | 171.0 | 29 | 0.0012 - 0.22 | 0.9995 | 1 |
| ***N*$^{\alpha}$-Acetyl-L-ornithine** | **AcOrn** | **$[^2H_4]$-ACC** | 345.1 | 171.0 | 37 | 0.0085 - 0.84 | 0.9995 | 1 |
| ***N*-Acetylputrescine** | **AcPut** | **$[^2H_4]$-Put** | 301.1 | 171.0 | 37 | 0.00085 - 0.84 | 0.9996 | 1 |
| **Agmatine** | **Agm** | **$[^2H_8]$-Agm** | 301.1 | 171.0 | 29 | 0.12 - 2.5 | 0.9997 | 1 |
| **L-Arginine** | **Arg** | **$[^2H_7]$-Arg** | 345.1 | 171.0 | 37 | 0.0057 - 5.6 | 0.9998 | 1 |
| **Cadaverine** | **Cad** | **$[^2H_4]$-Put** | 222.1 | 171.0 | 17 | 0.00045 - 0.045 | 0.9996 | 2 |
| **L-Citrulline** | **Cit** | **$[^2H_6]$-Cit** | 346.1 | 171.0 | 29 | 0.023 - 11 | 0.9997 | 1 |
| **Homospermidine** | **Hspd** | **$[^2H_8]$-Spm** | 335.6 | 171.0 | 29 | 0.00096 - 0.38 | 0.9992 | 3 |
| **L-Methionine** | **Met** | **$[^{13}C_5][^{15}N_1]$-Met** | 320.1 | 171.0 | 29 | 0.032 - 7.8 | 0.9990 | 1 |
| **L-Ornithine** | **Orn** | **$[^2H_6]$-Orn** | 473.1 | 171.0 | 37 | 0.28 - 9.8 | 0.9997 | 2 |
| **Putrescine** | **Put** | **$[^2H_4]$-Put** | 215.1 | 171.0 | 17 | 0.031 - 3.0 | 0.9988 | 2 |
| **Spermidine** | **Spd** | **$[^2H_8]$-Spd** | 328.6 | 171.0 | 29 | 0.079 - 1.5 | 0.9988 | 3 |
| **Spermine** | **Spm** | **$[^2H_8]$-Spm** | 295.2 | 171.0 | 17 | 0.057 - 2.2 | 0.9997 | 4 |
| **Thermospermine** | **Tspm** | **$[^2H_8]$-Spm** | 295.2 | 171.0 | 17 | 0.0045 - 1.1 | 0.9993 | 4 |
| | **$[^2H_4]$-ACC** | - | 276.1 | 171.0 | 29 | - | - | 1 |
| | **$[^2H_8]$-Agm** | - | 309.1 | 171.0 | 29 | - | - | 1 |
| | **$[^2H_7]$-Arg** | - | 352.1 | 171.0 | 37 | - | - | 1 |
| | **$[^2H_6]$-Cit** | - | 350.1 | 171.0 | 29 | - | - | 1 |
| | **$[^{13}C_5][^{15}N_1]$-Met** | - | 326.1 | 171.0 | 29 | - | - | 1 |
| | **$[^2H_6]$-Orn** | - | 479.2 | 171.0 | 37 | - | - | 2 |
| | **$[^2H_4]$-Put** | - | 217.1 | 171.0 | 17 | - | - | 2 |
| | **$[^2H_8]$-Spd** | - | 331.6 | 171.0 | 29 | - | - | 3 |
| | **$[^2H_8]$-Spm** | - | 297.8 | 171.0 | 17 | - | - | 4 |

Linear range and $R^2$ was not calculated for internal standards. Derivatization multiplicity indicates how many compatible amino groups have been derivatized.
*m/z* – mass-to-charge ratio, $R^2$ – coefficient of determination, eV – electron volts.

**Supplementary Table S2**: Summary of method validation parameters (for detailed description see chapter 2.3 Metabolite extraction and LC-MS/MS instrumentation).

| Analyte | Spiked content (pmol) | Average Method Accuracy (%bias) | Average Method Precision (%RSD) | Average Process Efficiency (%) | Average Matrix Effect (%) | Average Method Recovery (%) |
|---|---|---|---|---|---|---|
| **ACC** | 0.2 / 0.5 / 1 / 5 | 11.0 | 5.1 | 9.4 | 10.6 | 88.3 |
| **AcOrn** | 30 / 75 / 150 / 750 | 8.7 | 5.8 | 17.8 | 17.5 | 102.2 |
| **AcPut** | 3 / 7.5 / 15 / 75 | 14.5 | 1.4 | 32.1 | 32.8 | 96.9 |
| **Agm** | 280 / 700 | 13.8 | 13.9 | 7.6 | 39.7 | 110.2 |
| **Arg** | 400 / 1 000 / 2 000 | 3.9 | 4.6 | 27.9 | 248.2 | 100.5 |
| **Cad** | 0.04 / 0.1 / 0.2 | 12.2 | 3.0 | 53.0 | 44.6 | 119.5 |
| **Cit** | 800 / 2 000 / 4 000 | 8.0 | 4.6 | 35.2 | 15.2 | 210.7 |
| **Hspd** | 0.85 / 1.7 | 8.1 | 5.5 | 169.6 | 217.0 | 79.3 |
| **Met** | 28 / 70 / 140 / 700 | 10.4 | 2.0 | 34.7 | 32.4 | 106.4 |
| **Orn** | 700 / 1 750 | 5.5 | 6.4 | 24.5 | 29.8 | 78.6 |
| **Put** | 11 / 27.5 / 55 / 275 | 7.7 | 5.2 | 80.5 | 89.2 | 87.1 |
| **Spd** | 28 / 70 / 140 | 6.5 | 7.2 | 32.4 | 32.2 | 100.8 |
| **Spm** | 20 / 50 / 100 / 500 | 6.0 | 3.7 | 27.8 | 30.5 | 84.8 |
| **Tspm** | 4 / 10 / 20 | 11.6 | 4.6 | 65.9 | 55.3 | 113.4 |

Samples of *A*. *thaliana* (10 mg FW) were spiked with authentic standards as indicated in the table. Values for average method accuracy are expressed as %bias, values for average method precision are expressed as %RSD; n=4; FW = fresh weight; RSD, relative standard deviation. ACC, 1-aminocyclopropane-1-carboxylic acid; AcOrn, $N^{\alpha}$-acetyl-L-ornithine; AcPut, *N*-acetylputrescine; Agm, agmatine; Arg, L-arginine; Cad, cadaverine; Cit, L-citrulline; Hspd, homospermidine; Met, methionine; Orn, L-ornithine; Put, putrescine; Spd, spermidine; Spm, spermine; Tspm, thermospermine.

**Supplementary Table S3**: Autosampler and interday precision of studied polyamines and related compounds, expressed as %RSD.

| Analyte | Autosampler stability (n=3, %RSD) | | Interday precision (n=4; %RSD) |
|---|---|---|---|
| | Low | High | |
| **ACC** | 7.20 | 2.90 | 2.17 |
| **AcOrn** | 1.42 | 2.14 | 2.27 |
| **AcPut** | 1.45 | 6.84 | 1.03 |
| **Agm** | 7.82 | 1.00 | 14.61 |
| **Arg** | 4.58 | 10.08 | 3.82 |
| **Cad** | 8.68 | 2.13 | 2.46 |
| **Cit** | 9.53 | 2.49 | 3.22 |
| **Hspd** | 7.66 | 0.45 | 1.83 |
| **Met** | 3.36 | 2.50 | 1.84 |
| **Orn** | 3.05 | 0.10 | 1.76 |
| **Put** | 2.16 | 2.46 | 2.53 |
| **Spd** | 2.28 | 6.48 | 1.45 |
| **Spm** | 4.66 | 8.74 | 4.09 |
| **Tspm** | 4.66 | 2.64 | 12.71 |

Autosampler stability was examined by analyzing analyte standards at high (0.000005625 pmol ACC; 0.016875 pmol AcOrn; 0.0016875 pmol AcPut; 1.575 pmol Agm; 0.5625 pmol Arg; 0.00001125 pmol Cad; 0.1125 pmol Cit; 0.0019125 pmol Hspd; 0.0315 pmol Met; 0.0039375 pmol Orn; 0.061875 pmol Put; 0.1575 pmol Spd; 0.1125 pmol Spm; 0.01125 pmol Tspm) and low (0.225 pmol ACC; 33.75 pmol AcOrn; 3.375 pmol AcPut; 3.9375 pmol Agm; 5.625 pmol Arg; 0.045 pmol Cad; 11.25 pmol Cit; 0.3825 pmol Hspd; 31.5 pmol Met; 9.84375 pmol Orn; 12.375 pmol Put; 31.5 pmol Spd; 22.5 pmol Spm; 4.5 pmol Tspm) concentrations. Constant amount of IS mixture was added to each sample and measured on three consecutive days. Standards were kept in autosampler (6 °C) for the whole experiment duration. Interday precision was evaluated using four replicates of plant samples containing IS in the same intervals and conditions as for autosampler stability. RSD, relative standard deviation; IS, internal standards. ACC, 1-aminocyclopropane-1-carboxylic acid; AcOrn, $N^{\alpha}$-acetyl-L-ornithine; AcPut, *N*-acetylputrescine; Agm, agmatine; Arg, L-arginine; Cad, cadaverine; Cit, L-Citrulline; Hspd, homospermidine; Met, methionine; Orn, L-ornithine; Put, putrescine; Spd, spermidine; Spm, spermine; Tspm, thermospermine.

**Supplementary Table S4**: Examined analyte levels in seedlings of *A. thaliana* and *S. lycopersicum* under stress conditions and after treatment with aminoguanidine and L-norvaline.

| | | | | ACC | Cad | Hspd | Met | AcOrn | Put | AcPut | Spd | Spm | Tspm | Agm | Arg | Cit | Orn |
|---|---|---|---|---|---|---|---|---|---|---|---|---|---|---|---|---|---|
| | | | | pmol/g ± SD | pmol/g ± SD | pmol/mg ± SD | pmol/mg ± SD | pmol/mg ± SD | pmol/mg ± SD | pmol/mg ± SD | pmol/mg ± SD | pmol/mg ± SD | pmol/mg ± SD | pmol/mg ± SD | pmol/mg ± SD | pmol/mg ± SD | pmol/mg ± SD |
| Arabidopsis | Stress | | Control | 272.78 ± 24.62 | 20.37 ± 2.10 | 0.95 ± 0.12 | 13.97 ± 1.23 | 19.05 ± 3.11 | 64.49 ± 2.70 | 2.99 ± 0.08 | 165.16 ± 15.04 | 5.90 ± 0.56 | 2.59 ± 0.18 | 730.25 ± 22.13 | 941.94 ± 85.40 | 383.91 ± 38.43 | 620.46 ± 110.43 |
| | | Salinity | 10 mM | 147.12 ± 12.02 | 20.55 ± 10.96 | 0.92 ± 0.05 | 11.68 ± 0.37 | 7.65 ± 0.70 | 49.36 ± 1.63 | 2.25 ± 0.03 | 133.85 ± 10.43 | 5.10 ± 0.14 | 1.93 ± 0.05 | 586.36 ± 32.46 | 641.15 ± 55.29 | 226.84 ± 6.24 | 367.70 ± 15.49 |
| | | | 150 mM | 775.30 ± 27.18 | 7.03 ± 3.74 | 0.52 ± 0.02 | 12.27 ± 0.49 | 26.89 ± 3.24 | 41.87 ± 1.40 | 2.41 ± 0.22 | 102.01 ± 1.49 | 9.40 ± 0.58 | 0.43 ± 0.03 | 511.56 ± 18.21 | 354.78 ± 54.86 | 305.21 ± 18.91 | 140.63 ± 7.36 |
| | | | 300 mM | 215.30 ± 10.17 | ND | 0.16 ± 0.01 | 4.85 ± 0.12 | 6.64 ± 0.47 | 9.45 ± 0.37 | 0.37 ± 0.04 | 18.05 ± 1.16 | 2.20 ± 0.14 | 0.17 ± 0.01 | 135.29 ± 14.48 | 144.33 ± 24.79 | 40.26 ± 2.76 | 37.98 ± 6.40 |
| | | Drough | -0.5 MPa | 297.05 ± 29.48 | 43.14 ± 4.78 | 0.77 ± 0.06 | 64.48 ± 7.15 | 61.47 ± 4.76 | 157.27 ± 8.31 | 9.60 ± 1.61 | 221.52 ± 24.49 | 9.61 ± 1.04 | 0.78 ± 0.08 | 1983.65 ± 367.22 | 1637.90 ± 235.07 | 285.97 ± 37.74 | 406.83 ± 61.20 |
| | | | -0.7 MPa | 594.55 ± 91.36 | 54.79 ± 3.47 | 0.80 ± 0.08 | 100.45 ± 5.08 | 48.60 ± 10.64 | 181.83 ± 8.90 | 11.21 ± 2.02 | 259.95 ± 6.80 | 10.63 ± 0.66 | 0.85 ± 0.07 | 2808.65 ± 181.68 | 1894.60 ± 176.65 | 443.86 ± 32.55 | 520.34 ± 75.56 |
| | | | -1.2 MPa | 1083.96 ± 95.47 | 65.12 ± 7.76 | 0.91 ± 0.06 | 104.74 ± 11.10 | 38.71 ± 0.93 | 205.20 ± 16.59 | 13.36 ± 2.23 | 301.62 ± 8.96 | 12.88 ± 0.21 | 1.18 ± 0.12 | 3153.41 ± 273.43 | 1906.83 ± 205.18 | 311.34 ± 44.50 | 305.41 ± 21.73 |
| | Inhibitors | AG 48 h | Control | 166.68 ± 23.99 | 14.03 ± 3.80 | 1.14 ± 0.12 | 23.93 ± 2.06 | 5.71 ± 0.87 | 103.69 ± 7.05 | 2.58 ± 0.25 | 244.49 ± 30.70 | 6.01 ± 0.65 | 2.43 ± 0.27 | 854.67 ± 61.16 | 1031.30 ± 103.96 | 514.48 ± 32.48 | 119.79 ± 17.09 |
| | | | 0.1 mM | 267.98 ± 12.75 | 13.69 ± 1.37 | 1.11 ± 0.07 | 23.64 ± 1.35 | 9.96 ± 1.45 | 115.89 ± 1.41 | 2.50 ± 0.09 | 269.06 ± 12.19 | 6.52 ± 0.37 | 2.65 ± 0.15 | 944.40 ± 103.06 | 1043.67 ± 79.97 | 646.85 ± 33.79 | 188.58 ± 26.85 |
| | | | 1 mM | 281.78 ± 29.44 | 15.64 ± 3.40 | 1.02 ± 0.11 | 30.19 ± 1.77 | 7.32 ± 0.41 | 144.09 ± 6.59 | 3.25 ± 0.21 | 283.16 ± 23.22 | 6.73 ± 0.39 | 2.44 ± 0.16 | 1100.61 ± 135.68 | 925.63 ± 31.04 | 718.77 ± 30.09 | 139.73 ± 11.32 |
| | | | 10 mM | 3096.05 ± 655.59 | 36.51 ± 3.19 | 0.67 ± 0.03 | 24.03 ± 0.61 | 2.44 ± 0.55 | 353.43 ± 17.00 | 9.03 ± 1.07 | 458.08 ± 21.89 | 6.42 ± 0.28 | 1.76 ± 0.09 | 449.22 ± 53.26 | 1633.29 ± 66.62 | 432.82 ± 12.10 | 17.91 ± 1.96 |
| | | AG 96 h | Control | 97.73 ± 22.56 | 2.24 ± 0.94 | 0.89 ± 0.11 | 19.05 ± 1.35 | 3.85 ± 0.59 | 103.84 ± 5.37 | 3.16 ± 0.19 | 197.89 ± 22.90 | 6.67 ± 0.98 | 2.47 ± 0.41 | 818.70 ± 83.59 | 393.81 ± 47.68 | 294.70 ± 15.05 | 78.99 ± 10.49 |
| | | | 0.1 mM | 97.89 ± 13.90 | 2.20 ± 0.66 | 0.97 ± 0.08 | 21.24 ± 1.41 | 6.24 ± 1.26 | 112.64 ± 3.27 | 3.23 ± 0.16 | 234.62 ± 19.24 | 7.88 ± 0.65 | 2.92 ± 0.22 | 1002.41 ± 82.71 | 530.88 ± 65.64 | 378.57 ± 12.22 | 134.97 ± 26.34 |
| | | | 1 mM | 102.08 ± 16.30 | 3.46 ± 0.92 | 0.90 ± 0.06 | 21.59 ± 1.36 | 6.94 ± 1.13 | 128.03 ± 5.21 | 4.46 ± 0.43 | 224.96 ± 25.15 | 7.84 ± 0.30 | 2.65 ± 0.12 | 707.94 ± 99.72 | 550.20 ± 48.86 | 404.96 ± 18.58 | 152.07 ± 29.71 |
| | | | 10 mM | 6951.67 ± 1384.96 | 23.73 ± 2.98 | 0.52 ± 0.09 | 27.23 ± 0.95 | 7.11 ± 0.96 | 485.69 ± 61.84 | 25.53 ± 2.77 | 466.41 ± 45.94 | 7.29 ± 0.27 | 1.49 ± 0.08 | 320.20 ± 30.79 | 1377.01 ± 103.10 | 453.30 ± 38.70 | 23.01 ± 4.47 |
| | | Nor 48 h | Control | 355.28 ± 43.47 | 70.00 ± 14.90 | 0.42 ± 0.03 | 15.32 ± 0.78 | 8.02 ± 1.33 | 33.37 ± 0.35 | 1.21 ± 0.03 | 273.05 ± 11.10 | 6.67 ± 0.35 | 1.87 ± 0.07 | 473.73 ± 97.19 | 1755.25 ± 232.19 | 281.81 ± 20.10 | 55.33 ± 9.79 |
| | | | 0.1 mM | 369.82 ± 24.49 | 60.71 ± 8.44 | 0.75 ± 0.03 | 29.24 ± 0.88 | 10.49 ± 1.15 | 44.65 ± 2.35 | 1.44 ± 0.06 | 278.67 ± 10.14 | 7.25 ± 0.50 | 1.92 ± 0.11 | 511.08 ± 68.70 | 1949.84 ± 113.57 | 470.64 ± 9.29 | 71.63 ± 9.72 |
| | | | 1 mM | 853.79 ± 145.57 | 78.39 ± 5.89 | 0.84 ± 0.06 | 60.95 ± 3.08 | 23.46 ± 2.38 | 89.44 ± 6.81 | 2.66 ± 0.38 | 290.78 ± 14.99 | 8.15 ± 0.58 | 1.74 ± 0.22 | 472.76 ± 41.24 | 1995.55 ± 273.47 | 805.12 ± 21.43 | 209.76 ± 4.98 |
| | | | 10 mM | 4729.97 ± 547.52 | 140.65 ± 30.38 | 1.39 ± 0.19 | 95.37 ± 4.09 | 53.76 ± 9.87 | 169.27 ± 12.76 | 6.19 ± 1.17 | 280.80 ± 23.16 | 7.61 ± 0.61 | 1.98 ± 0.28 | 839.12 ± 72.91 | 2165.40 ± 588.58 | 1406.02 ± 58.30 | 757.15 ± 18.07 |
| | | Nor 96 h | Control | 74.61 ± 8.38 | 60.50 ± 14.63 | 0.44 ± 0.02 | 13.66 ± 1.07 | 3.85 ± 0.67 | 37.93 ± 1.99 | 1.53 ± 0.17 | 229.37 ± 35.58 | 6.17 ± 0.42 | 1.73 ± 0.07 | 412.67 ± 43.13 | 973.65 ± 105.87 | 185.67 ± 10.60 | 28.87 ± 5.21 |
| | | | 0.1 mM | 95.88 ± 8.17 | 43.79 ± 2.36 | 0.43 ± 0.04 | 15.25 ± 0.68 | 4.86 ± 0.51 | 33.68 ± 2.49 | 1.45 ± 0.26 | 210.94 ± 14.67 | 6.32 ± 0.37 | 1.68 ± 0.11 | 495.06 ± 56.36 | 1471.95 ± 93.64 | 247.26 ± 7.26 | 43.59 ± 6.11 |
| | | | 1 mM | 362.28 ± 47.67 | 58.89 ± 4.51 | 0.73 ± 0.02 | 53.54 ± 7.17 | 55.09 ± 5.42 | 101.77 ± 11.57 | 6.00 ± 0.60 | 324.99 ± 29.25 | 9.78 ± 0.81 | 1.69 ± 0.05 | 721.02 ± 122.00 | 2010.05 ± 227.99 | 844.85 ± 97.60 | 155.49 ± 27.85 |
| | | | 10 mM | 3664.40 ± 720.25 | 119.62 ± 33.01 | 1.37 ± 0.23 | 88.39 ± 2.46 | 50.19 ± 10.05 | 232.79 ± 19.69 | 12.02 ± 1.46 | 395.68 ± 74.26 | 9.76 ± 1.64 | 2.40 ± 0.36 | 1056.97 ± 133.61 | 2395.93 ± 383.37 | 1783.15 ± 136.18 | 837.52 ± 155.04 |

| | | | | ACC | Cad | Hspd | Met | AcOrn | Put | AcPut | Spd | Spm | Tspm | Agm | Arg | Cit | Orn |
|---|---|---|---|---|---|---|---|---|---|---|---|---|---|---|---|---|---|
| | | | | pmol/g ± SD | pmol/g ± SD | pmol/mg ± SD | pmol/mg ± SD | pmol/mg ± SD | pmol/mg ± SD | pmol/mg ± SD | pmol/mg ± SD | pmol/mg ± SD | pmol/mg ± SD | pmol/mg ± SD | pmol/mg ± SD | pmol/mg ± SD | pmol/mg ± SD |
| Tomato | Stress | | Control | 63.13 ± 13.69 | 217.72 ± 30.65 | 5.84 ± 0.90 | 16.94 ± 0.96 | 2.79 ± 0.32 | 122.72 ± 6.84 | 7.72 ± 0.70 | 149.20 ± 19.20 | 4.76 ± 0.30 | 2.12 ± 0.23 | 1.05 ± 0.15 | 606.41 ± 108.32 | 285.03 ± 34.01 | 130.10 ± 15.04 |
| | | Salinity | 10 mM | 57.70 ± 7.21 | 200.71 ± 27.12 | 5.34 ± 0.74 | 20.36 ± 2.00 | 2.86 ± 0.31 | 123.04 ± 13.58 | 7.29 ± 0.30 | 159.72 ± 7.56 | 5.51 ± 0.47 | 2.59 ± 0.40 | 0.83 ± 0.14 | 645.58 ± 102.36 | 307.92 ± 21.94 | 138.04 ± 14.49 |
| | | | 150 mM | 47.60 ± 5.85 | 173.30 ± 19.33 | 4.31 ± 0.60 | 16.05 ± 1.76 | 2.22 ± 0.08 | 88.81 ± 9.20 | 7.46 ± 1.05 | 174.04 ± 21.94 | 7.93 ± 0.67 | 2.15 ± 0.25 | 1.10 ± 0.23 | 474.06 ± 89.16 | 268.09 ± 42.92 | 127.39 ± 26.32 |
| | | | 300 mM | 4,907.61 ± 1328.46 | 136.07 ± 12.82 | 3.10 ± 0.15 | 24.11 ± 2.09 | 3.77 ± 0.63 | 59.06 ± 4.79 | 6.94 ± 0.58 | 147.22 ± 3.89 | 9.49 ± 0.55 | 1.03 ± 0.13 | 1.24 ± 0.11 | 983.05 ± 107.06 | 633.60 ± 45.65 | 130.45 ± 6.64 |
| | | Drough | -0.5 MPa | 87.19 ± 6.83 | 376.00 ± 44.09 | 6.60 ± 1.10 | 26.49 ± 2.89 | 5.14 ± 0.45 | 278.39 ± 48.82 | 22.47 ± 4.67 | 266.05 ± 45.18 | 4.71 ± 0.79 | 1.60 ± 0.20 | 5.06 ± 0.86 | 1457.44 ± 119.62 | 373.84 ± 68.64 | 212.19 ± 32.04 |
| | | | -0.7 MPa | 103.65 ± 22.08 | 464.97 ± 68.74 | 5.91 ± 0.33 | 35.45 ± 3.30 | 7.58 ± 1.02 | 365.06 ± 31.50 | 28.76 ± 2.59 | 237.12 ± 19.79 | 3.97 ± 0.70 | 1.94 ± 0.35 | 4.74 ± 1.64 | 1820.04 ± 302.29 | 422.70 ± 65.84 | 253.11 ± 43.59 |
| | | | -1.2 MPa | 237.17 ± 71.57 | 396.80 ± 87.35 | 5.38 ± 0.28 | 37.27 ± 1.84 | 12.76 ± 1.20 | 576.75 ± 85.68 | 47.83 ± 6.70 | 263.13 ± 52.80 | 3.61 ± 0.57 | 2.04 ± 0.13 | 6.79 ± 1.67 | 2350.71 ± 305.10 | 411.57 ± 51.81 | 216.81 ± 68.00 |
| | Inhibitors | AG 48 h | Control | 62.36 ± 10.48 | 288.17 ± 18.28 | 4.92 ± 0.49 | 17.85 ± 0.97 | 5.54 ± 0.67 | 154.65 ± 9.78 | 11.80 ± 1.27 | 203.46 ± 17.06 | 3.79 ± 0.32 | 1.08 ± 0.10 | 1.88 ± 0.25 | 963.86 ± 104.16 | 238.23 ± 20.66 | 136.75 ± 14.47 |
| | | | 0.1 mM | 62.10 ± 8.79 | 280.15 ± 20.45 | 4.57 ± 0.24 | 17.86 ± 1.10 | 6.62 ± 0.25 | 160.25 ± 9.26 | 13.05 ± 1.69 | 217.04 ± 20.04 | 3.77 ± 0.22 | 1.36 ± 0.20 | 1.31 ± 0.21 | 1001.34 ± 62.37 | 299.73 ± 21.32 | 194.79 ± 41.53 |
| | | | 1 mM | 33.69 ± 5.44 | 281.48 ± 22.29 | 4.04 ± 0.34 | 17.50 ± 0.37 | 6.49 ± 0.53 | 150.28 ± 11.39 | 14.96 ± 1.28 | 188.40 ± 3.11 | 4.95 ± 0.70 | 1.23 ± 0.20 | 1.19 ± 0.15 | 1010.63 ± 90.55 | 299.28 ± 36.56 | 220.39 ± 44.17 |
| | | | 10 mM | 28.17 ± 9.61 | 365.98 ± 57.74 | 4.36 ± 0.46 | 29.01 ± 3.15 | 4.78 ± 0.69 | 198.35 ± 24.14 | 18.67 ± 2.39 | 187.13 ± 38.28 | 3.79 ± 0.45 | 0.99 ± 0.15 | 1.42 ± 0.53 | 928.98 ± 228.09 | 202.31 ± 30.52 | 224.29 ± 39.87 |
| | | AG 96 h | Control | 77.73 ± 9.16 | 463.22 ± 36.93 | 6.01 ± 0.53 | 15.77 ± 0.74 | 8.16 ± 0.61 | 183.15 ± 13.56 | 19.99 ± 1.47 | 223.45 ± 18.82 | 3.81 ± 0.23 | 1.18 ± 0.05 | 3.75 ± 0.94 | 1105.93 ± 226.12 | 246.14 ± 38.12 | 182.15 ± 23.61 |
| | | | 0.1 mM | 98.21 ± 1.94 | 479.70 ± 32.46 | 5.00 ± 0.83 | 15.30 ± 0.26 | 6.91 ± 0.37 | 183.09 ± 19.81 | 20.77 ± 4.24 | 212.62 ± 13.29 | 4.07 ± 0.18 | 1.29 ± 0.07 | 2.77 ± 0.90 | 884.24 ± 130.00 | 290.03 ± 26.14 | 137.03 ± 22.50 |
| | | | 1 mM | 87.68 ± 15.37 | 366.40 ± 34.49 | 3.71 ± 0.71 | 13.26 ± 0.93 | 5.82 ± 0.73 | 177.87 ± 14.06 | 20.50 ± 1.90 | 208.80 ± 12.52 | 3.86 ± 0.23 | 1.67 ± 0.15 | 1.70 ± 0.52 | 988.08 ± 132.20 | 226.43 ± 35.13 | 139.46 ± 25.72 |
| | | | 10 mM | 23.62 ± 2.45 | 518.95 ± 87.78 | 2.69 ± 0.28 | 23.76 ± 1.25 | 6.11 ± 0.85 | 258.16 ± 17.69 | 31.17 ± 3.42 | 197.69 ± 11.09 | 4.03 ± 0.32 | 1.67 ± 0.10 | 2.15 ± 0.58 | 1012.22 ± 59.27 | 162.46 ± 24.90 | 140.17 ± 25.92 |
| | | Nor 48 h | Control | 33.67 ± 6.12 | 244.86 ± 25.82 | 5.55 ± 0.46 | 17.65 ± 1.81 | 7.58 ± 1.03 | 132.84 ± 11.56 | 12.42 ± 1.66 | 190.99 ± 23.85 | 5.05 ± 0.53 | 1.09 ± 0.13 | 2.69 ± 0.41 | 641.61 ± 46.23 | 216.30 ± 32.44 | 140.64 ± 12.20 |
| | | | 0.1 mM | 66.29 ± 6.62 | 252.81 ± 18.26 | 5.07 ± 0.39 | 20.17 ± 1.45 | 7.93 ± 1.13 | 157.87 ± 17.73 | 12.37 ± 1.18 | 195.43 ± 20.55 | 4.81 ± 0.37 | 1.01 ± 0.14 | 1.65 ± 0.26 | 871.69 ± 84.35 | 254.03 ± 39.65 | 204.30 ± 30.24 |
| | | | 1 mM | 94.11 ± 8.77 | 295.80 ± 24.58 | 7.15 ± 0.76 | 33.20 ± 1.91 | 10.09 ± 0.51 | 164.77 ± 14.72 | 14.57 ± 1.74 | 190.15 ± 18.26 | 5.22 ± 0.22 | 1.07 ± 0.09 | 1.97 ± 0.45 | 850.13 ± 133.42 | 526.38 ± 32.97 | 374.18 ± 29.28 |
| | | | 10 mM | 132.86 ± 41.21 | 224.54 ± 27.92 | 6.52 ± 1.05 | 85.67 ± 7.30 | 16.75 ± 2.15 | 206.82 ± 22.92 | 15.55 ± 2.61 | 210.02 ± 40.51 | 3.80 ± 0.43 | 0.93 ± 0.10 | 1.55 ± 0.24 | 1074.59 ± 138.06 | 1051.01 ± 132.70 | 2055.70 ± 332.49 |
| | | Nor 96 h | Control | 98.95 ± 19.58 | 356.60 ± 18.46 | 7.58 ± 0.74 | 15.35 ± 0.58 | 7.60 ± 0.65 | 158.10 ± 3.93 | 21.56 ± 1.88 | 262.38 ± 22.29 | 4.57 ± 0.20 | 1.77 ± 0.16 | 4.35 ± 0.94 | 601.31 ± 107.24 | 236.01 ± 17.01 | 109.02 ± 18.89 |
| | | | 0.1 mM | 155.36 ± 19.95 | 373.59 ± 13.84 | 8.42 ± 1.53 | 17.24 ± 0.82 | 8.85 ± 0.42 | 168.19 ± 15.18 | 18.05 ± 1.34 | 269.66 ± 42.28 | 4.47 ± 0.11 | 1.64 ± 0.12 | 4.63 ± 1.12 | 695.94 ± 79.04 | 297.81 ± 27.54 | 156.79 ± 31.25 |
| | | | 1 mM | 153.01 ± 35.15 | 392.36 ± 15.06 | 7.04 ± 0.75 | 33.13 ± 2.12 | 10.83 ± 1.11 | 200.71 ± 16.34 | 24.28 ± 3.54 | 257.99 ± 51.40 | 4.00 ± 0.15 | 1.74 ± 0.12 | 3.57 ± 0.86 | 720.08 ± 65.41 | 517.14 ± 45.42 | 435.91 ± 78.34 |
| | | | 10 mM | 156.57 ± 26.60 | 270.08 ± 35.62 | 4.42 ± 0.53 | 130.30 ± 5.74 | 13.95 ± 2.76 | 300.60 ± 38.04 | 28.07 ± 6.63 | 218.36 ± 16.02 | 2.86 ± 0.25 | 1.19 ± 0.07 | 5.47 ± 1.48 | 1323.96 ± 217.81 | 1630.90 ± 124.92 | 3271.00 ± 383.22 |

Results expressed in pmol/g FW (ACC, Cad) and in pmol/mg for remaining analytes. Values expressed as means ± SD, n = 4-5. AG, aminoguanidine; Nor, L-norvaline; FW, fresh weight; ND, not detected.

ACC, 1-aminocyclopropane-1-carboxylic acid; AcOrn, $N^{\alpha}$-acetyl-L-ornithine; AcPut, *N*-acetylputrescine; Agm, agmatine; Arg, L-arginine; Cad, cadaverine; Cit, L-citrulline; Hspd, homospermidine; Met, methionine; Orn, L-ornithine; Put, putrescine; Spd, spermidine; Spm, spermine; Tspm, thermospermine.

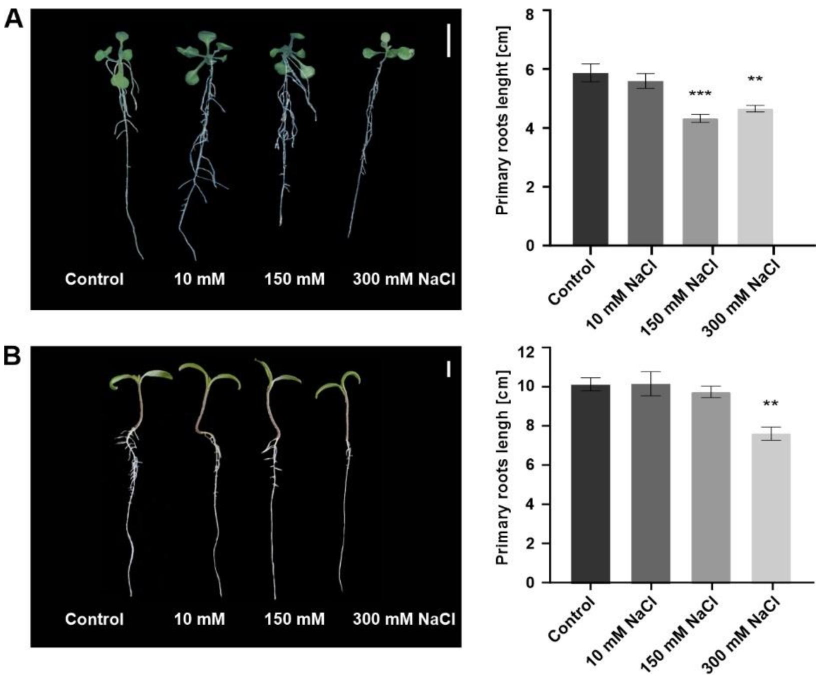


**Supplementary Fig. S1**: Phenotypes of *A*. *thaliana* and *S*. *lycopersicum* after 48 h of salinity stress treatment.

Phenotype of 12 DAG old *A*. *thaliana* (A) and 6 DAG old S. *lycopersicum* (B) seedlings subjected to salinity stress at three concentration levels (10 mM; 150 mM and 300 mM) for 48 h. Comparison of seedlings primary root length (right) measured using ImageJ software. Values calculated from 4-8 seedlings. Error bars represent SE, the number of asterisks indicates significance level (* = $p<0.05$; ** = $p<0.01$; *** = $p<0.001$) in comparison to control, non-stressed plants, using Welch's ANOVA Test. Scale bar = 10 mm; SE, standard error.

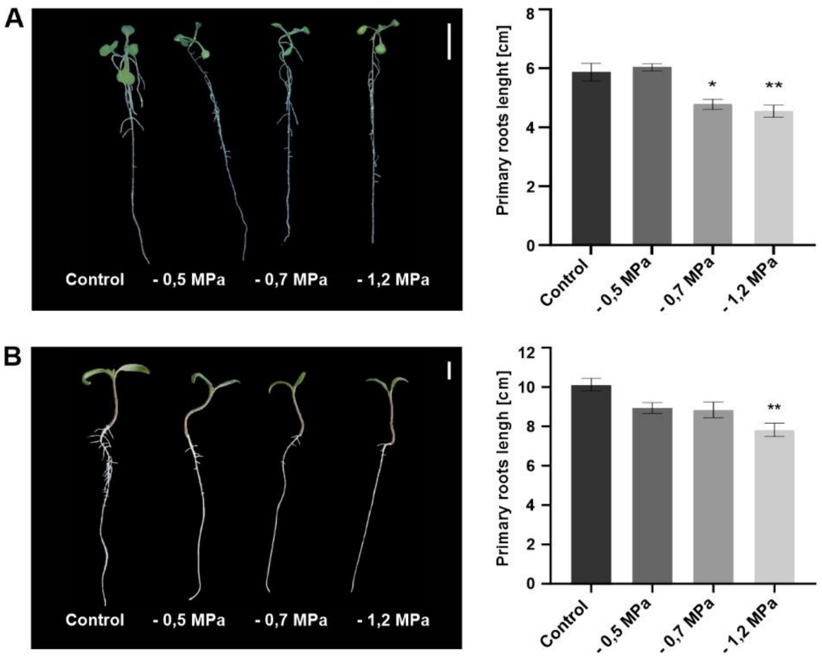


**Supplementary Fig. S2**: Phenotypes of *A*. *thaliana* and *S*. *lycopersicum* after 48 h of drought stress treatment.

Phenotype of 12 DAG old *A*. *thaliana* (A) and 6 DAG old S. *lycopersicum* (B) seedlings subjected to drought stress at three concentration levels (-0.5 MPa; -0.7 MPa and -1.2 MPa) for 48 h. Comparison of seedlings primary root length (right) measured using ImageJ software. Values calculated from 4-8 seedlings. Error bars represent SE, the number of asterisks indicates significance level (* = p<0.05; ** = p<0.01; *** = p<0.001) in comparison to control, non-stressed plants, using Welch's ANOVA Test. Scale bar = 10 mm; SE, standard error.

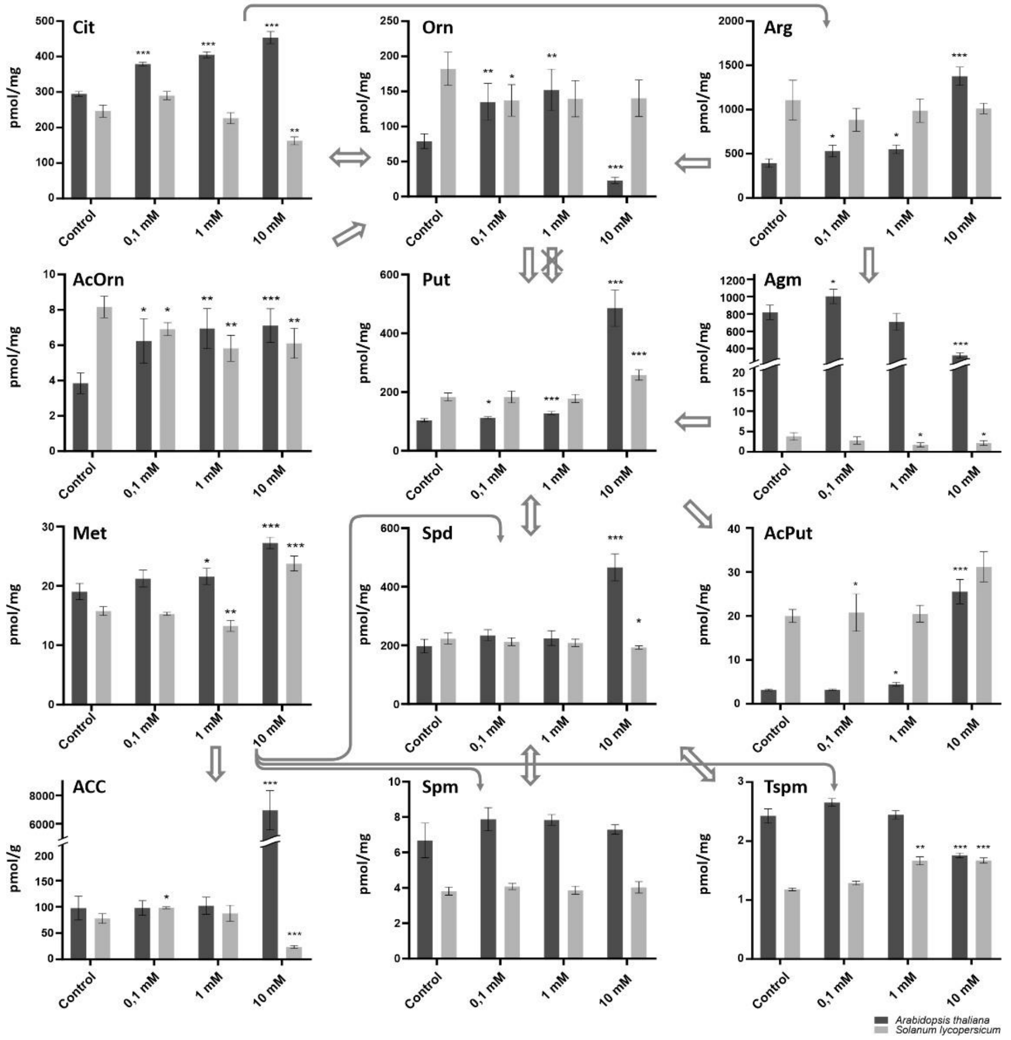


**Supplementary Fig. S3:** Polyamine and related compounds levels in aminoguanidine treated *A*. *thaliana* and *S*. *lycopersicum* after 96 h of treatment.

Metabolite profiles of polyamines and related compounds of 9 DAG old *A*. *thaliana* (dark bars) and 8 DAG old *S*. *lycopersicum* (light bars) seedlings treated with aminoguanidine (AG) for 96 h at three concentration levels, 0.1 mM, 1 mM and 10 mM. The major polyamines Put, Spd, Spm and Tspm content is depicted along with related amino acids and biogenic amines (Cit, Arg, Orn, Agm), metabolite forms (AcOrn, AcPut) and Yang cycle metabolites (Met, ACC). Metabolite concentrations calculated from 4-5 technical replicates are expressed in pmol/g FW (ACC) and in pmol/mg FW for all other analytes. Error bars represent SD, the number of asterisks indicates significance level (* = p<0.05; ** = p<0.01; *** = p<0.001) in comparison to control, non-treated plants, using Welch's ANOVA Test. The arrows represent metabolic fluxes between the compounds, crossed out arrow indicates absence of the responsible enzyme in *Arabidopsis*, based on Lou et al., 2020.

ACC, 1-aminocyclopropane-1-carboxylic acid; AcOrn, $N^{\alpha}$-acetyl-L-ornithine; AcPut, *N*-acetylputrescine; Agm, agmatine; Arg, L-arginine; Cit, L-citrulline; Met, methionine; Orn, L-ornithine; Put, putrescine; Spd, spermidine; Spm, spermine; Tspm, thermospermine; DAG, days after germination; FW, fresh weight; SD, standard deviation.

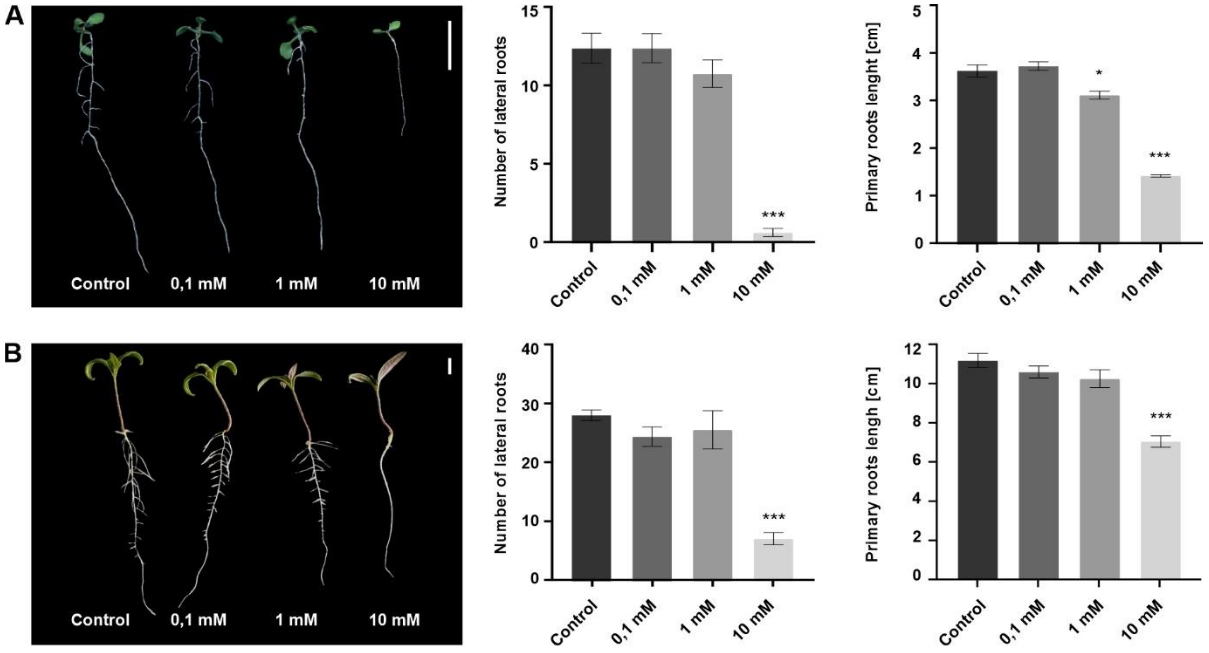


**Supplementary Fig. S4:** Phenotypes of aminoguanidine treated *A*. *thaliana* and *S*. *lycopersicum* after 96 h of treatment.

Phenotype of 9 DAG old *A*. *thaliana* (A) and 8 DAG old *S*. *lycopersicum* (B) seedlings treated with aminoguanidine (AG) for 96 h at three concentration levels, 0.1 mM, 1 mM and 10 mM. Comparison of number of lateral roots of seedlings (central) and the length of their primary roots (right) measured using ImageJ software. Values calculated from 4-8 seedlings. Error bars represent SE, the number of asterisks indicates significance level (* = p<0.05; ** = p<0.01; *** = p<0.001) in comparison to control, non-treated plants, using Welch's ANOVA Test. Scale bar = 10 mm; SE, standard error.

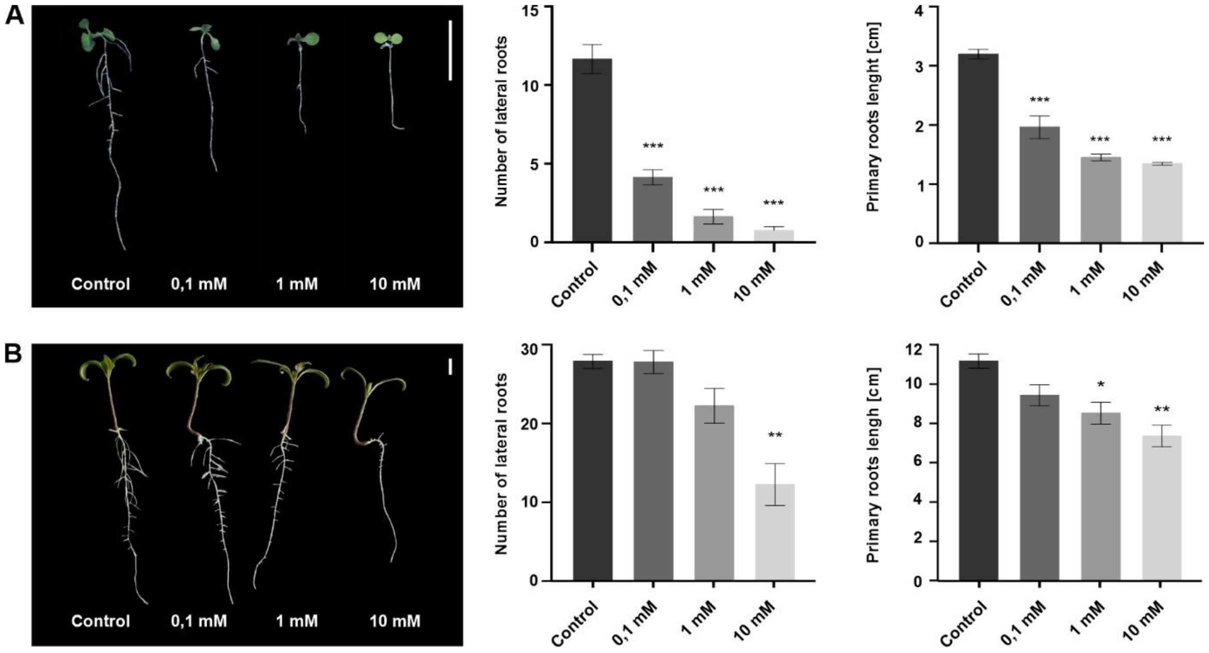


**Supplementary Fig. S5:** Phenotypes of L-norvaline treated *A. thaliana* and *S. lycopersicum* after 96 h of treatment.

Phenotype of 9 DAG old *A. thaliana* (A) and 8 DAG old *S. lycopersicum* (B) seedlings treated with L-norvaline (Nor) for 96 h at three concentration levels, 0.1 mM, 1 mM and 10 mM. Comparison of number of lateral roots of seedlings (central) and the length of their primary roots (right) measured using ImageJ software. Values calculated from 4-8 seedlings. Error bars represent SE, the number of asterisks indicates significance level (* = $p<0.05$; ** = $p<0.01$; *** = $p<0.001$) in comparison to control, non-treated plants, using Welch's ANOVA Test. Scale bar = 10 mm; SE, standard error.

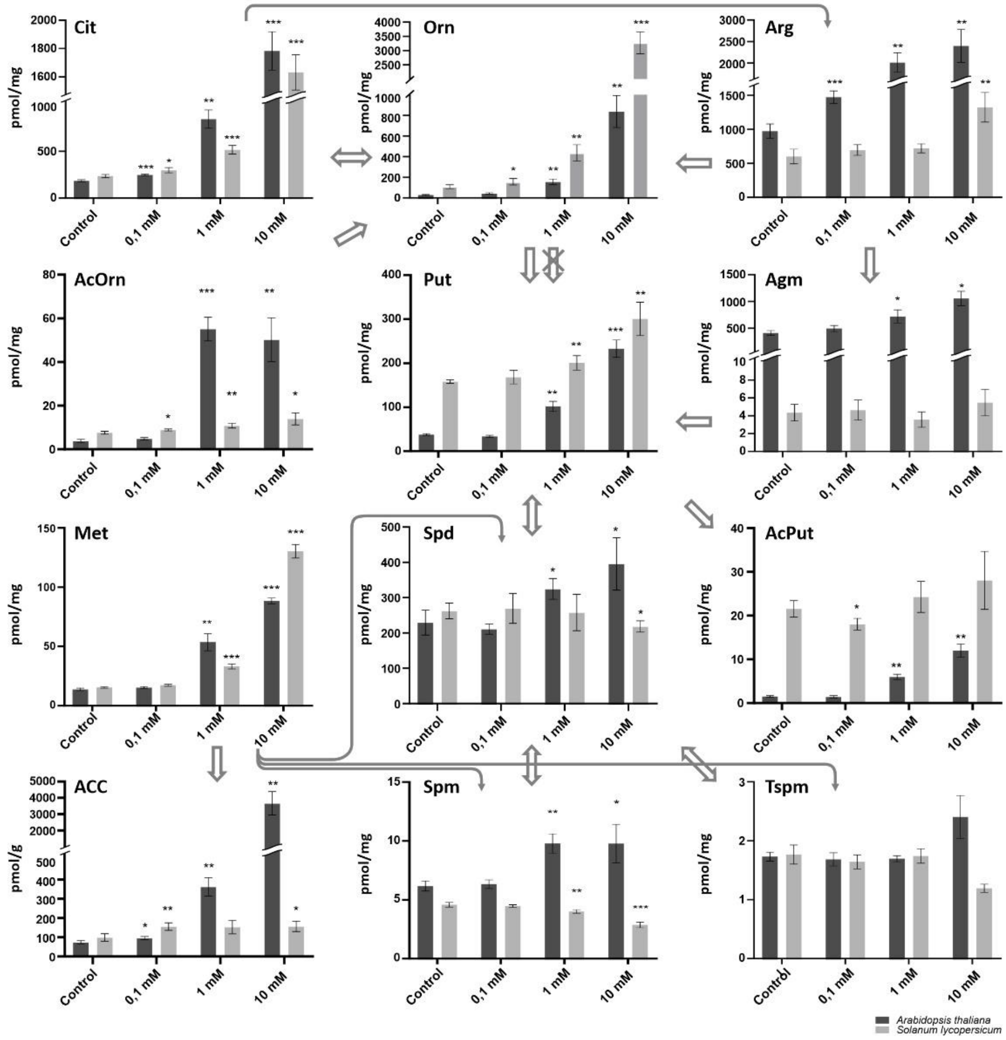


**Supplementary Fig. S6:** Polyamine and related compounds levels in L-norvaline treated *A. thaliana* and *S. lycopersicum* after 96 h of treatment.

Metabolite profiles of polyamines and related compounds of 9 DAG old *A. thaliana* (dark bars) and 8 DAG old *S. lycopersicum* (light bars) seedlings treated with L-norvaline (Nor) for 96 h at three concentration levels, 0.1 mM, 1 mM and 10 mM. The major polyamines Put, Spd, Spm and Tspm content is depicted along with related amino acids and biogenic amines (Cit, Arg, Orn, Agm), metabolite forms (AcOrn, AcPut) and Yang cycle metabolites (Met, ACC). Metabolite concentrations calculated from 4-5 technical replicates are expressed in pmol/g FW (ACC) and in pmol/mg FW for all other analytes. Error bars represent SD, the number of asterisks indicates significance level (* = p<0.05; ** = p<0.01; *** = p<0.001) in comparison to control, non-treated plants, using Welch's ANOVA Test. The arrows represent metabolic fluxes between the compounds, crossed out arrow indicates absence of the responsible enzyme in *Arabidopsis*, based on Lou et al., 2020.

ACC, 1-aminocyclopropane-1-carboxylic acid; AcOrn, Nα-acetyl-L-ornithine; AcPut, N-acetylputrescine; Agm, agmatine; Arg, L-arginine; Cit, L-citrulline; Met, methionine; Orn, L-ornithine; Put, putrescine; Spd, spermidine; Spm, spermine; Tspm, thermospermine; DAG, days after germination; FW, fresh weight; SD, standard deviation.
